# Co-transcriptional splicing changes combine with reduced productive transcription initiation for cold-induced repression of *FLC*

**DOI:** 10.1101/2025.06.20.660682

**Authors:** Robert Maple, Govind Menon, Susan Duncan, Maria Sindalovskaya, Hongchun Yang, Rebecca Bloomer, Martin Howard, Caroline Dean

**Author notes:** These authors contributed equally to this work (Robert Maple, Govind Menon).

## Abstract

The Arabidopsis floral repressor locus *FLC* is epigenetically silenced during winter cold to align flowering with spring. During weeks of cold exposure, *FLC* transcription is progressively reduced both by transcriptional repression mediated by *FLC* antisense transcription, and epigenetic silencing implemented through a Polycomb-mediated epigenetic switch. In the warm, *FLC* is transcriptionally repressed by coordinated changes in transcription initiation and RNA PolII speed in a mechanism involving proximal termination. Whether similar mechanisms contribute to the cold-induced *FLC* transcriptional repression is unknown. Here, we combine mathematical modelling and transcription profiling to investigate *FLC* transcriptional changes during the cold. We find different dynamics of spliced and unspliced transcripts during cold exposure with only a small change in PolII speed. We also show that, unlike short-term cold, long-term cold temperatures drive an increase in splicing rates while simultaneously reducing productive transcription at *FLC*. This process is influenced by antisense *COOLAIR* transcription but does not rely on proximal *COOLAIR* termination. Cold-induced transcriptional repression of *FLC* thus involves a decoupling of changes in productive transcription initiation from PolII speed and rates of co-transcriptional splicing, a different mechanism to that repressing *FLC* in the warm.

## INTRODUCTION

Transcription is a complex process requiring numerous co-transcriptional steps that act co-ordinately to generate a mature transcript. Transcription initiation, promoter proximal pause release, PolII processivity, PolII elongation, splicing and termination are all opportunities for regulation of RNA levels [1]. Transcription can be further influenced by a multiplicity of mechanisms including nucleosome remodelling altering promoter accessibility, transcription factor binding modulating PolII recruitment, as well as changes in intrinsic PolII activity and in the chromatin landscape through which PolII transcribes [2]. In addition to this regulation, quality control pathways can direct transcripts for RNA turnover [3], whilst positive feedbacks connect the different co-transcriptional processes adding to the complexity [4–6]. Only a fraction of initiated transcription events result in a mature transcript making transcription appear highly inefficient; this multilayered control, however, enables a high degree of regulation.

One locus where transcriptional dynamics have been studied in detail is the *Arabidopsis* floral repressor *FLOWERING LOCUS C (FLC).* Quantitative expression of this gene contributes to life history choice in a wide range of plants [7]; a high level determines an overwintering requirement and one generation per year. A constitutively low level of expression underpins a rapid-cycling strategy, enabling multiple generations a year where environmental conditions allow. The difference in expression is set during early development in the embryo, promoted by the *FLC* transcriptional activator FRIGIDA (FRI) [8], and repressed by activity of the autonomous flowering pathway including FCA [9]. FRIGIDA and FCA function antagonistically in a transcription-coupled repression mechanism that links proximal transcription termination with the histone methylation status of the local chromatin, determining the timing of the subsequent switch to a fully silenced Polycomb state [10, 11]. Thus, a balance between FCA and FRI quantitatively regulates *FLC* expression in a graded manner in the warm, before switching to a Polycomb silenced digital OFF state. In the warm, we have previously found a >20-fold difference in transcription initiation between highly-transcribed and Polycomb silenced states, with ∼8-10 fold changes in elongation rates (PolII speed) [12]. The FCA-mediated transcription-coupled repression mechanism in the warm affects both *FLC* sense and antisense transcription, with feedback mechanisms that reinforce both strands, potentially explaining why *FLC* is so sensitive to disruption of these co-transcriptional regulators. These aspects are summarised schematically in Fig. S1.

High transcription of *FLC* is progressively downregulated during winter in the process of vernalization, aligning flowering with spring. The cold-induced reduction in *FLC* expression involves both transcriptional repression and epigenetic silencing, where we reserve “silencing” to mean epigenetically stable inhibition of gene expression, with “repression” referring to non-epigenetic down-regulation. These two mechanisms can operate independently, are capable of responding on different timescales, and work in parallel to reduce *FLC* expression [13]. *COOLAIR* transcription is induced in the cold and is mutually exclusive to *FLC* expression at a single allele [14, 15]. *COOLAIR* also contributes to cold-induced *FLC* transcriptional repression through sequestration of the activator FRI into dynamic biomolecular condensates, located away from the *FLC* locus [16]. However, a full understanding of how cold temperatures achieve *FLC* transcriptional repression is still to be achieved. The complex interactions between the different phases of the transcription cycle and difficulties in interpreting transcriptional data complicate this investigation. For example, distinguishing between changes to transcription initiation and elongation can be difficult, as both change the levels of nascent transcripts but in opposite directions. Changes to the dynamics of different transcripts, such as spliced and unspliced, can also be explained in multiple ways and not just through a changed splicing rate, e.g., through different RNA stabilities. Such mechanisms can only be disentangled and understood through integration of computational modelling and experiments.

Here, we develop a predictive mathematical model parameterised with experimental data to analyse *FLC* transcriptional repression in the cold. We show that repression involves coordinated changes at multiple transcriptional steps, is not the same as the repression mechanism in the warm and does not involve proximal transcriptional termination. Interestingly, we find that a reduced productive transcription initiation rate, with only a small contribution from reduced PolII speed, coexists with an enhanced co-transcriptional splicing rate. Overall, these data support a mechanism where PolII coupling to co-transcriptional splicing is modulated by temperature. Cold temperature reduces productive *FLC* transcriptional initiation but paradoxically results in a higher rate of splicing.

## RESULTS

### Measurement of spliced to unspliced *FLC* ratio reveals slower shut-down dynamics for spliced RNA during cold treatment

Cold treatment leads to a progressive reduction in levels of *FLC* mRNA over weeks of vernalization (Fig. 1A). This reduction is mediated by a combination of transcriptional repression and epigenetic silencing, through concurrent targeting of the *FLC* locus by two pathways – antisense mediated transcriptional repression and Polycomb Repressive Complex 2 (PRC2) mediated epigenetic silencing [13]. We previously showed that in Col*FRI*, the antisense repression pathway is capable of responding to temperature changes both on a fast timescale of hours to days and on a slow timescale over multiple weeks [13, 17, 18], while the PRC2 pathway only responds on a slow timescale of weeks [13]. Under constant 5°C treatment, both pathways respond on a slow timescale, jointly contributing to reducing *FLC* transcriptional output. To examine the resulting changes to *FLC* transcription under constant 5°C conditions, we started by measuring *FLC* mRNA levels in non-vernalized Col*FRI* plants (NV) and after cold treatments of 2, 4, 6 and 8 weeks. For *FLC* mRNA (the spliced RNA), the forward primer anneals from nucleotide number 4265 relative to the Transcription Start Site (TSS) and spans spliced exons 4 and 5 while the reverse primer is entirely within the last exon (exon 7). We also measured levels of *FLC* RNA where the intron 2-exon 3 junction is yet to be spliced, which we term unspliced *FLC* RNA. *FLC* unspliced primers span from nucleotide numbers 3966 to 4136 relative to the TSS of *FLC*. The amplicon they generate starts within intron 2 and ends within intron 3. If only productive transcription initiation is changing under cold treatment, we would expect spliced and unspliced *FLC,* measured using the above primers, to exhibit similar quantitative changes. However, the reduction in spliced *FLC* (*FLC* mRNA) relative to NV was consistently smaller than the corresponding fold reduction in unspliced *FLC*, throughout the cold time course (Fig. 1A). Interestingly, the difference in the rate of reduction is most pronounced in the initial phase of cold (NV to 2W), after which spliced and unspliced transcript levels change at similar rates (Fig. 1A). The slower reduction in spliced relative to unspliced *FLC* is a consistent feature of the cold-induced reduction in *FLC* transcriptional output in all our previously published datasets [15, 16, 18–20]. To examine whether the cold-induced difference between spliced and unspliced *FLC* persists in the post-cold warm conditions, we examined plants 7 days after each cold-treatment (Fig. 1A). Interestingly, the difference is mostly absent in the post-cold warm, indicating that this is a cold-specific effect.

**Figure 1.**
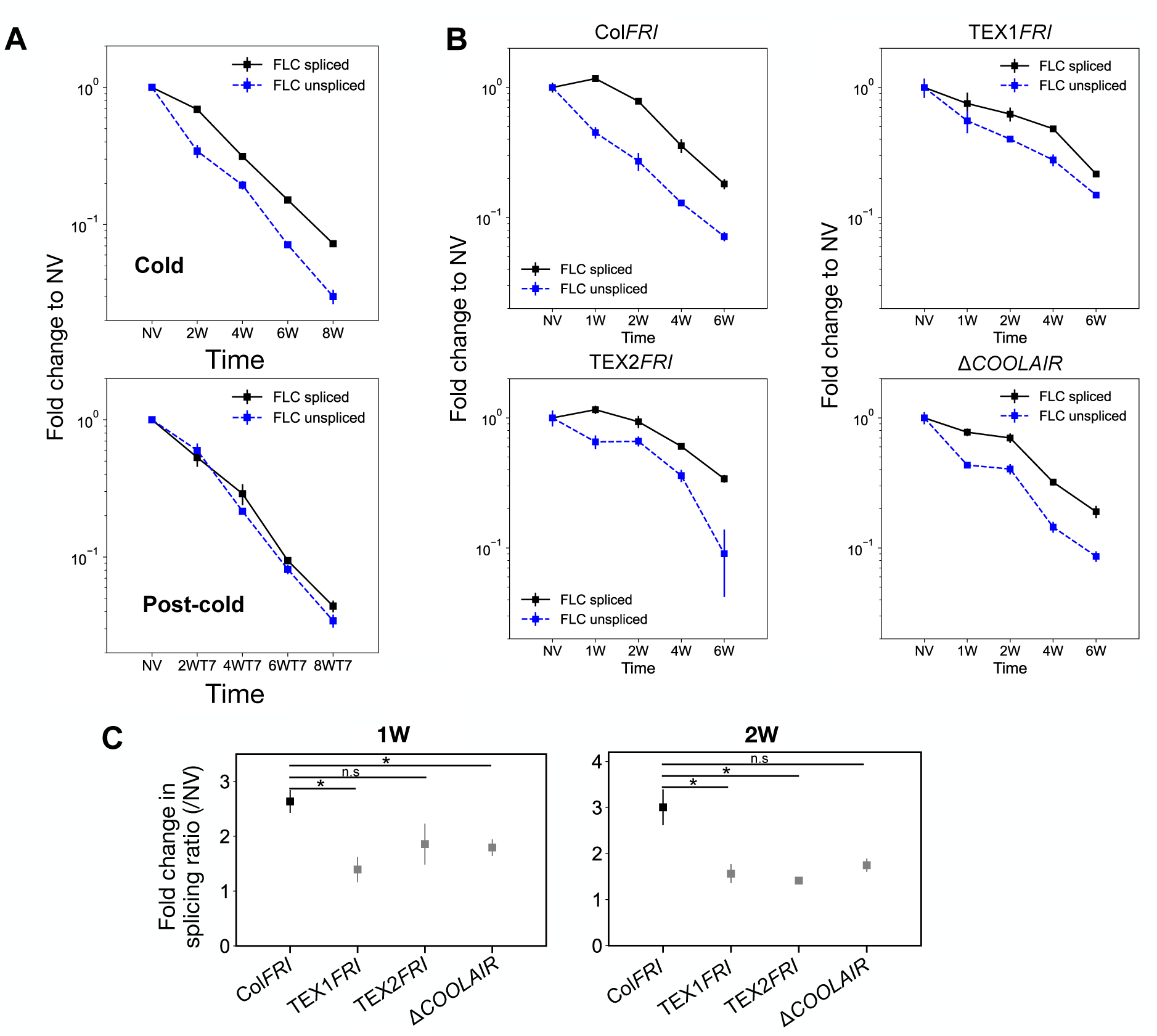
Cold induced changes in processing of *FLC* sense transcription. Relative expression of *FLC* mRNA and unspliced *FLC* over a vernalization time course in (A) Wild-type (WT) (Col*FRI*) and (B) WT and *COOLAIR* mutant lines. Values are normalized to the housekeeping gene *PP2A* and to the non-vernalized (NV) levels, error bars represent SEM (n=3). (C) Fold change relative to NV of splicing ratio (spliced *FLC* RNA: unspliced *FLC* RNA) over a vernalization time course. Values are normalized to the housekeeping gene *PP2A* and Col*FRI* starting levels, error bars represent SEM (n=3 biological replicates). Since the trend was the same across all three *COOLAIR* defective mutants, a one-tailed Student’s t-test was used to compare each mutant to Col*FRI*. The Bonferroni correction was used to adjust the significance level from α = 0.05 to α = 0.0167 (for three comparisons). (*) indicates p < 0.0167; n.s indicates no significance (p ≥ 0.0167).

The differences between spliced and unspliced dynamics were most pronounced in the first two weeks of cold exposure, when *COOLAIR* is being strongly up-regulated. We therefore examined whether *COOLAIR* transcription affected the *FLC* spliced and unspliced RNA dynamics by comparing three different antisense defective mutants compared to the wild-type over a 6-week cold time course, using data from Nielsen et al., 2024 [13]. The differences between the spliced and unspliced RNA dynamics were reduced compared to the wild-type (Fig. 1B,C) suggesting that some aspects of *COOLAIR* transcription could influence the mechanism underlying the different dynamics.

### Slow cold-induced repression of *FLC* mRNA does not involve changed mRNA stability

Previous work in *S. cerevisiae* showed that increased antisense transcription at certain gene loci can lead to enhanced stability of mRNA transcribed from these loci [21]. If the *FLC* locus follows this paradigm, we would expect long-term cold to enhance *FLC* mRNA lifetime, since antisense *COOLAIR* transcription is upregulated at the *FLC* locus during long-term cold [13, 18, 19]. Such a change in stability could explain the difference in dynamics of spliced versus unspliced *FLC* RNA. To directly test the altered stability hypothesis, we measured *FLC* mRNA turnover after transcriptional arrest with the transcriptional inhibitor Actinomycin D (ActD) (Fig. 2A,B). Previous measurements of *FLC* half-life were performed under non-vernalized conditions [9, 22] or have estimated RNA turnover relative to a housekeeping gene by qPCR [19]. To accurately measure absolute mRNA abundance, we carried out smFISH probing *FLC* spliced mRNA on non-vernalized (NV) and 2-week vernalized (2W) seedlings after ActD treatment.

**Figure 2.**
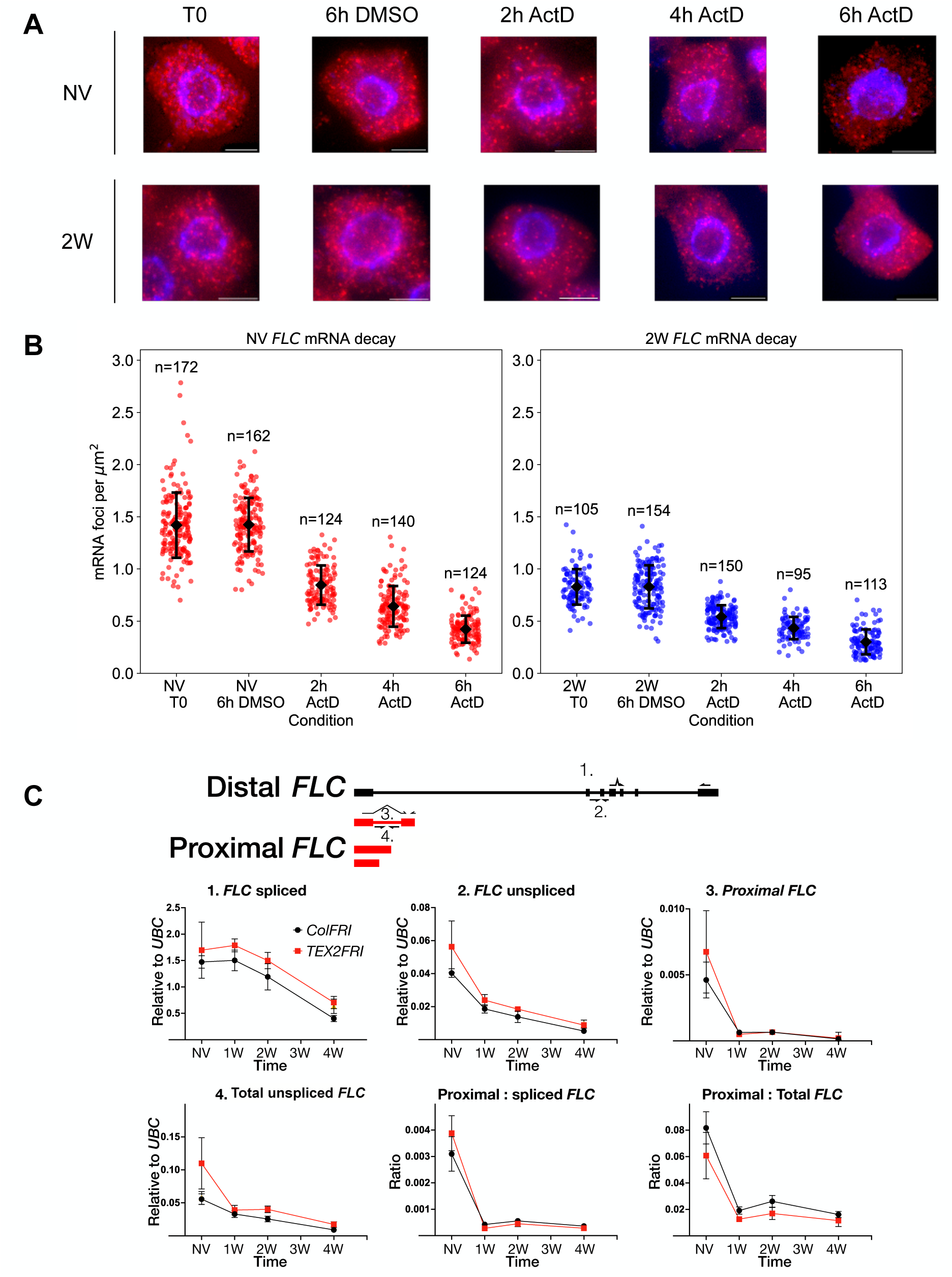
*FLC* mRNA stability and *FLC* proximal termination compared between warm and cold treated plants. (A) Representative smFISH images showing spatial distribution of individual *FLC* mRNA foci (red) in root meristem cells. Non-vernalized (NV) and 2-week cold treated plants (2W) were exposed to the transcriptional inhibitor Actinomycin D (ActD) for 0, 2, 4 or 6 hours. Nuclear staining from DAPI is shown in blue. Scale bars = 5 µm. (B) *FLC* mRNA quantified relative to cell area. Dots indicate individual cell mRNA counts relative to cell area (μm^2^). NV samples (red), cold samples (blue). Error bars in (B) indicate mean +/- standard deviation. Number of cells in each sample indicated. (C) Transcript dynamics of *FLC* isoforms in the cold for Col*FRI* and TEX2 *FRI*. Isoform schematics (top) with canonical *FLC* (black) and proximal isoforms (red). Transcript dynamics for spliced (1), unspliced (2), proximal spliced (3), and total unspliced *FLC* (4) over a vernalization time course. Isoform schematic shows primer measurement locations for (1-4). Proximal *FLC* isoforms also shown relative to spliced and total unspliced *FLC* transcript levels, respectively (bottom). Values are normalized to the housekeeping gene *UBC* and *FLC* NV levels (error bars represent SEM (n=4)).

Starting *FLC* mRNA levels were reduced after 2-week cold treatment by ∼0.6-fold going from 1.4 foci per µm^2^ per cell to 0.8 foci per µm^2^ per cell, consistent with previous findings of *FLC* transcriptional repression (Fig. 2B). Treatment with ActD led to a progressive reduction in the number of mRNA foci for *FLC* as the mRNA is degraded (Fig. 2A, B). We fit an exponential decay separately to the NV and 2-week data to estimate *FLC* decay rate constants as 0.18 ± 0.009 hr^-1^ (ordinary least squares estimate: mean ± standard error) for warm conditions and 0.16 ± 0.009 hr^-1^ for cold conditions giving estimated half-lives of ∼3.9 hours and ∼4.4 hours respectively. These half-lives are shorter, but comparable to previous estimates of *FLC* mRNA half-life of ∼6 hours [22] and ∼5 hours [9]. The difference in decay rate between NV and 2-week data was not statistically significant (ANCOVA: t-test for difference in decay rate constant between conditions: p=0.12). The similar half-life of *FLC* mRNA in warm and cold conditions indicates that *FLC* mRNA stability is not significantly changed after 2 weeks of cold treatment. As *COOLAIR* transcription is known to increase significantly by 2 weeks [14, 19], it is therefore unlikely that *COOLAIR* transcription significantly changes *FLC* mRNA stability. Furthermore, the unchanged half-life implies that altered mRNA turnover cannot be a major contributor to the difference in dynamics between spliced and unspliced transcripts. Thus, the long-term cold-induced dynamics of spliced and unspliced *FLC* may reflect an increased rate of splicing, potentially affected by *COOLAIR* transcription.

### A simple mathematical analysis can quantify the splicing rate changes

The increase in splicing rate – here interpreted as the rate at which the intron is cleaved at the 3’ splice site and the exons are ligated - can be quantified (for the intron 2-exon 3 splice site being measured) using a simple ordinary differential equation model for the dynamics of *FLC* spliced and unspliced transcripts. The model assumes that all of the unspliced transcripts are eventually converted to spliced transcripts. See Supplementary Information for a description of the model. At steady state, the spliced to unspliced ratio is then equal to the ratio of the splicing rate to the decay rate of the spliced transcript. Since the decay rate of the spliced transcript is unchanged in cold versus warm conditions, the fold change in this ratio between the cold time points and NV is equal to the fold change in splicing rate between these conditions. For the data in Fig. 1A, this calculation gives a splicing rate fold change of 2.0 ± 0.2 at the 2W time point, while for the Col*FRI* data in Fig. 1B, it gives a fold change of 3.0 ± 0.4 at the same time point. The fold-change in the ratio continues to be significantly higher than 1 at the 4-, 6-, and 8-week timepoints (Fig. S2). These estimates suggest a clear increase in the splicing rate in the cold. The fold-change at the post-cold timepoints is closer to 1 in all cases (after 2, 4, 6, and 8 week cold treatments, Fig. S2), indicating that the increased splicing rate is a cold specific effect, and the splicing rate reverts to a lower value upon return to warm conditions.

### Cold-induced *FLC* repression does not involve enhanced proximal polyadenylation of sense transcription

Establishment of the transcriptionally repressed state of *FLC* in early development in the warm involves proximal polyadenylation that results in specific PolII-associated factors delivering a repressive chromatin environment [10, 11]. If the cold-induced transcriptional-repression involves a similar mechanism, we would expect to see an increase in proximal polyadenylation of *FLC* sense transcription under cold treatment.

To assess premature polyadenylation of *FLC* sense transcription in the cold we took advantage of a reported proximally polyadenylated, spliced isoform, (Fig. 2C) [8], to develop a qPCR assay to quantitatively measure proximal polyadenylation at that site. This proximal *FLC* isoform reduced faster than spliced or unspliced *FLC* over a 4 week cold timecourse (Fig. 2C). We also measured “total unspliced *FLC*” using a primer that captures all transcripts that are unspliced covering the proximal isoform. The amplicon covers nucleotides 612-698 relative to the TSS (see Supplemental Table 2). After 1 week of cold exposure, the ratio of the proximal *FLC* isoform to total unspliced *FLC* was reduced by ∼4-fold relative to NV, and this change was maintained over the rest of the cold exposure time course (Fig. 2C). Considering that antisense transcription can influence *FLC* shutdown dynamics (Fig. 2C), the *COOLAIR* defective *TEX2 FRI* was also included in the analysis (Fig. 2C). Both starting levels and cold dynamics for the proximally polyadenylated isoform were found to be similar between Col*FRI* and TEX2 *FRI*. Thus, cold-induced *FLC* repression correlates with reduced, rather than enhanced formation of this proximally polyadenylated isoform. Whether other proximal isoforms are affected, or polyadenylation-independent termination mechanisms are at play, will require further investigation.

### plaNETseq analysis indicates a cold-induced reduction in productive transcription initiation

To continue our investigation of which aspect of transcription was affected by cold we carried out plaNETseq to assay PolII position over *FLC* at single-nucleotide resolution (Fig. 3A). plaNETseq only maps the 3’ end of the nascent transcript, or the final nucleotide incorporated by the PolII active site, which can be used as a proxy for PolII position on the locus. To isolate all phosphorylated forms of engaged PolII in an unbiased way, we took advantage of a FLAG-tagged PolII complementation line crossed into the *ColFRI* background with an active *FLC* locus [23]. To further enhance sequencing of *FLC* transcripts we exploited an in-solution hybridisation capture method. After enrichment, the correlation remained high between replicates (Fig. S3). We also found the expected dynamics for the control genes *VIN3* (upregulated in cold conditions) and *ACT7* (remains constant) (Fig. S4).

**Figure 3.**
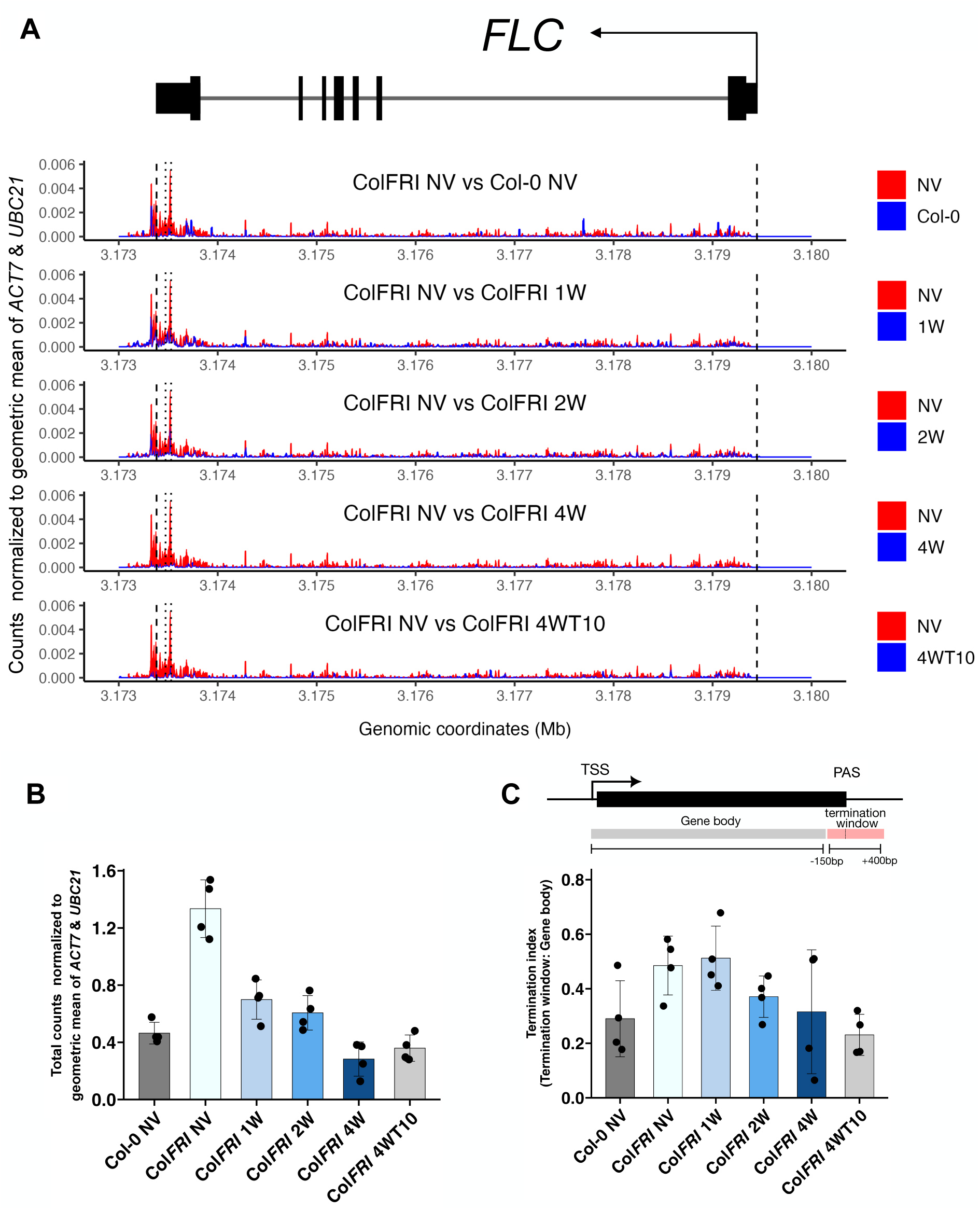
plaNETseq analysis of the FLC locus over a cold time course. (A) PolII distributions across FLC over the vernalization time series. Sense plaNETseq read signal from 4 replicates was normalized to the geometric mean of ACT7 and UBC21. Each panel shows a comparison between a cold timepoint and NV conditions in ColFRI or between Col-0 NV and ColFRI NV. Distributions are plotted as curves and the space between the curves is shaded blue where the cold timepoint or Col-0 NV shows higher Pol II signal and red where the ColFRI NV timepoint shows higher Pol II signal. FLC TSS and the end of 3’ UTR (Araport 11 annotation) are represented by black dashed lines and major poly-A sites are shown by black dotted lines. (B) Total plaNETseq counts over FLC over the cold timecourse. Mean counts of 4 biological replicates normalized to the geometric mean of ACT7 and UBC21. (C) PolII termination index (TI) at FLC over the vernalization time series decreases in the cold. TI was calculated by finding the ratio of counts around the first major polyA site (PAS) of FLC (-150 to +400 bp) relative to the remaining gene body counts. In (B) and (C), n=4 biological replicates at all timepoints. Error bars indicate mean +/- standard deviation in (B) and (C).

Like previous observations[12], the strongest relative enrichment of PolII in warm-grown plants was over the 3’ end of the locus, including over the major *FLC* polyadenylation sites (polyadenylation sites observed in our previous Quantseq analysis [10]) and beyond these sites, potentially associated with transcription termination (Fig. 3A). Enrichment over the transcription start site (TSS) and the gene body was lower. The overall pattern of low enrichment over the gene body and higher enrichment over the TTS did not change significantly through a cold exposure time course (Fig. S5) (1, 2, 4 weeks constant cold exposure). There were no locations within the gene body where an obvious drop in overall signal beyond this location could be observed. Hence these profiles did not suggest the presence of major early termination sites (agreeing with our analysis in the previous section).

We then examined PolII pausing across *FLC.* A PolII stalling site just within the 5’ end of introns had been detected in previous plaNETseq analyses when comparing plants given 3h or 12h of cold exposure, particularly in genes with long introns and with convergently transcribed antisense transcripts (CAS) [23]. The first intron of *FLC* is one of the longest introns in Arabidopsis and antisense transcription at *FLC* is well documented. However, only a mild relative enrichment of PolII was detected at the beginning of intron 1 (Fig. 3A). The profiles also did not show obvious cold-induced pause sites. We examined whether *FLC* has relative PolII enrichment at the VAL1 binding site within intron 1; a platform for co-ordinating transcriptional repression [24]. However, plaNETseq enrichment in the first intron of *FLC* was not significantly cold-induced.

We expected to see a reduction in overall PolII levels across *FLC* in the cold, consistent with the reduced transcriptional output observed in our time course experiments. This was indeed the case (Fig. 3B), with a progressive reduction in PolII levels over time in the cold. As expected from the well-studied epigenetic maintenance of *FLC* silencing in the post-cold, PolII levels did not recover to NV levels at the post-cold (4WT10) time point. We also analysed the *FLC* locus in the Col-0 genotype using plaNETseq. Consistent with the epigenetically silenced state of *FLC* in this genotype [11, 12], we observed significantly lower PolII levels in Col-0 seedlings compared to non-vernalized Col*FRI* (Fig. 3A,B). The fold changes in overall PolII levels (Fig. 3B) were broadly consistent with the expected fold reduction in spliced *FLC* RNA at the corresponding cold timepoints. While the fold changes in Fig. 3B relative to NV are larger than those in Fig. 1A, we note that these changes are variable between experiments. Larger changes in spliced *FLC* RNA at 2W and 4W, consistent with those in Fig. 3B have been observed in our previous datasets [18, 24]. The consistent fold changes in PolII levels and spliced *FLC* RNA, combined with the unchanged stability of spliced *FLC* RNA described above, indicates that the changes in overall PolII level are mainly caused by reduced transcription initiation, and there is no sharp decrease in PolII speed over *FLC* in the cold.

To quantify any overall change in PolII speed over the gene body, we examined the ratio of total reads in the polyadenylation and termination window relative to the total reads over the gene body (Fig. 3C – see figure caption for the definition of this window). Here we call this ratio the termination index. This ratio was reduced in the cold, indicating a shift in the relative residence time of PolII within these regions during transcription in different environmental conditions. The observed changes in Fig. 3C are consistent with two alternative scenarios (or a combination) – either PolII slows down slightly over the gene body while its speed remains unchanged over the termination window, or PolII speeds up slightly over the termination window while its speed over the gene body is unchanged. We favour a situation close to the first scenario involving a slow down over the gene body in cold conditions, since such a slowdown is consistent with predictions from our previous model describing co-transcriptional H3K36me3 histone modifications at *FLC* over the same cold time points [13]. Our model [13], which was based on the link between PolII residence time over a region and the probability of adding H3K36me3 in that region, was fit to H3K36me3 changes across *FLC* over a vernalization time course for Col*FRI* and *COOLAIR* defective mutants. The fitted model predicted a reduction in PolII speed across *FLC* in the cold (fold change of 0.6 at all cold timepoints) [13]. In this study, we used a simple mathematical model of PolII distribution over the locus to infer PolII speed changes from the plaNETseq profiles (see Supplementary Information for a description of this model). Based on this model, comparing the termination index (Fig. 3C) at the NV and 2-week time points gives an estimate of an 0.75 mean fold change in PolII elongation rate over the gene body, which is approximately consistent with our previous estimate. The termination index is essentially unchanged between NV and 1-week, indicating little change in the PolII elongation rate at this earlier timepoint. At the later, 4-week timepoint, the mean termination index reduces further, indicating a mean fold change of 0.63 in the PolII elongation rate. However, we note the high variability between replicates at this timepoint. Comparing the termination index at the post-cold 4WT10 timepoint to NV indicates a further reduction to 0.47, consistent with further PolII slowdown in an H3K27me3 spread state (Fig. 3C). The termination index at 4WT10 is comparable to its value for Col-0 seedlings, where *FLC* is fully covered by H3K27me3 (Fig. 3C) [11, 12].

Thus, in summary, the plaNETseq data supports the conclusion that the strongest effect of the early cold phase on sense *FLC* transcription is a change to productive transcriptional initiation. The large changes in either processivity or elongation rate (∼10 fold) that occur in silencing in the warm are not observed in the early cold phase [12].

### Coordinated changes in productive transcription initiation and PolII speed alone are insufficient to explain cold-induced changes in intron 1 total RNA

Analysis of total RNA changes alone (e.g., data in Fig. 1A) does not provide sufficient information to uniquely determine all the underlying transcriptional changes in the cold. Rates of initiation/transition to elongation, PolII speed, and splicing rates all contribute to different aspects of transcriptional dynamics in different ways. To examine these changes on the sense strand, we therefore performed a quantitative analysis of *FLC* intron 1 processing, using a mathematical model for steady-state RNA levels across an intron [12]. This modelling approach allows us to tease apart the contributions from the individual co-transcriptional processes and examine how they collectively generate the observed RNA dynamics. Previously, this approach was successfully applied to reveal that the underlying transcriptional changes between an active transcriptional state and a Polycomb-silenced state at *FLC* in the warm comprised coordinated reductions in transcription initiation and PolII speed [12].

To understand whether cold-induced transcriptional repression involved a similar coordinated change of initiation and PolII speed, we first analysed the total RNA fraction using the same model as in [12], comparing non-vernalized (NV) and vernalized conditions (after 2 weeks of vernalization, 2W). We measured RNA levels by qPCR at different locations by tiling primers across the intron. The model incorporates parameters capturing transcription initiation, PolII speed, intron processing rate, and 5’-3’ intron lariat degradation (Fig. 4A). Fitting the model to the qPCR measured total RNA levels across the intron, and constraining the model by known ranges of elongation rate and estimated basal splicing rate [12, 22], allowed us to examine which of these parameters are changed between NV and 2W and by how much. The measured levels show a clear increasing trend towards the 3’ end of the intron (Fig. 4B). Surprisingly, we found that a coordinated reduction in productive transcription initiation and PolII speed alone, as in [12], could not capture the changes between NV and 2W. Even the qualitative changes predicted by the model were in the wrong direction and therefore failed to match the data (Fig. S6). However, reduced initiation (fold change of 0.67±0.07) and an increased intron processing rate (fold change of 6.9±3.5), with the modest experimentally derived reduction in elongation from the previous section (0.8 fold) could produce good fits to the data (Fig. S7). To obtain these values for initiation and processing, we performed model parameter inference using a Bayesian approach [25], using a nested sampling algorithm [26] (see Supplementary Information and Table S1).

**Figure 4.**
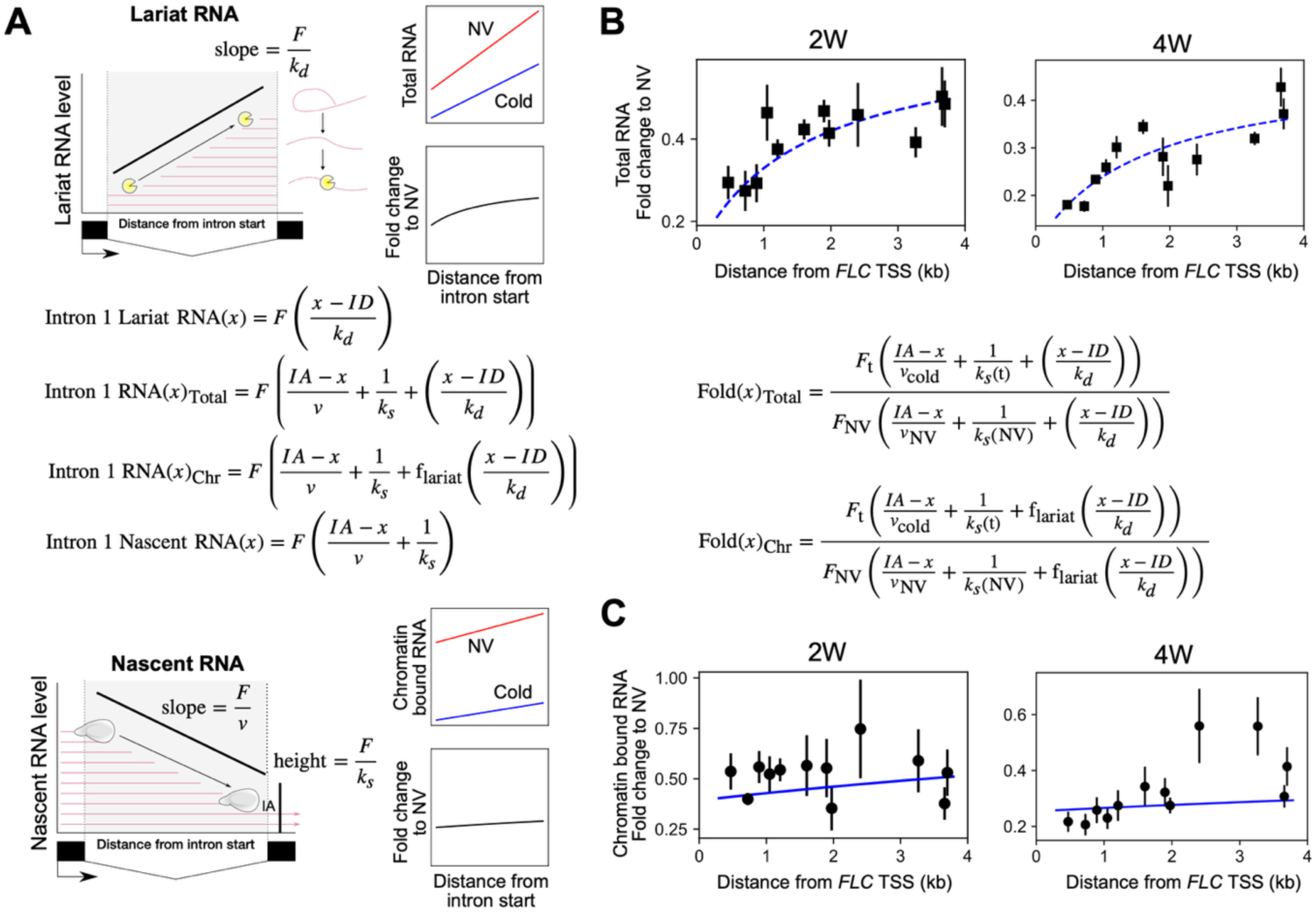
Mathematical model of RNA levels across *FLC* intron 1 predicts reduced productive transcription initiation and increased intron processing rate for *FLC* sense transcription in the cold. (A) Schematic representation of intron lariat and nascent RNA contribution to RNA levels across intron 1, and their relationship to productive transcription initiation, intron processing rate and lariat degradation rate. Top: schematic representation of lariat RNA profile: lariat turnover products generate a characteristic slope determined by the initiation rate *F* and 5’-3’ lariat turnover rate *k_d_*. Bottom: schematic representation of nascent RNA profile across intron 1 of *FLC*. The slope along the intron reflects the PolII density or a ratio of the initiation rate *F* to the PolII elongation speed *ν*. Signal level at the intron acceptor site *IA* corresponds to the ratio of the initiation rate to the intron processing rate *k_s_*. Model equations for the nascent and lariat RNA contributions are shown. From these we obtain model equations for the total and chromatin-bound RNA fractions by assuming that all of the lariat component contributes to the total RNA fraction, while only a fraction of the lariat component contributes to the chromatin-bound fraction (as captured by the parameter *f*_lariat_). (B,C) Model predictions indicated by blue lines – dashed for total RNA and solid for chromatin bound RNA. Experimental datapoints indicated by black squares (total RNA) and black circles (chromatin bound RNA). (B) Model fits for total RNA fold change profile across *FLC* intron 1 (relative to NV profile). Top left: 2W, Top right: 4W. Bottom left: Model equation for total fold change. The following model parameters were inferred using the intron 1 tiling data: fold change in productive initiation, fold change in intron processing rate, and the lariat contribution parameter *f*_lariat_. See Figs. S3-S6 for inferred values. Parameter inference was carried out using a nested sampling approach (see Supplementary Information). Model fits shown (dashed blue curves) are obtained using the mean values of the estimated posterior distribution for each of the parameters. (C) As for (B) but for chromatin-bound RNA fold change across *FLC* intron 1. Model equation for chromatin-bound fold change shown at top right. Model fits shown (full blue curves). Error bars in (B) and (C) represent SEM (n=3 biological replicates).

We note that the intron processing rate parameter in the intron 1 model represents a composite of a sequence of steps by which intronic RNA, which is part of the nascent transcript, is converted to an intron lariat subject to degradation (see Supplementary Information). It is thus not exactly equivalent to the splicing rate interpreted as the rate of cleavage at the 3’ splice site. Nevertheless, the inferred increase in this composite rate parameter for intron 1 is consistent with the increased rate of cleavage found above for intron 2, coupled with an overall increased rate of the other intron processing steps. This agreement indicates that an increased splicing rate in the cold is likely to apply to all *FLC* introns.

While a cold-induced change to PolII speed across intron 1 was not essential to generate good model fits, a slight slowdown in PolII is indicated by previous data [13], as well as by the plaNETseq data presented above. We therefore used the above experimentally derived elongation rate changes in our subsequent analysis. We next attempted to validate these predictions by using the estimated changes in initiation and intron processing rates from the total RNA fits to predict the profile of intron 1 nascent (chromatin-bound) RNA.

### Reduced productive transcription initiation and enhanced intron processing rate together explain changes to both intron 1 total and chromatin-bound RNA

We used the parameter values estimated from the total RNA profile to predict the chromatin-bound RNA profiles, using the corresponding model from Wu et al.[12]. This model assumes no contribution from the intron lariat to the chromatin-bound RNA fraction (Fig. 4C). We found that the best-fit model parameters from the total RNA analysis predicted a decreasing trend across the intron for chromatin-bound RNA. However, when we repeated the intron 1 tiling analysis for the chromatin-bound RNA fraction, the data did not show a decreasing trend across the intron (Fig. 4C). Even allowing for the fact that the fold change in *FLC* initiation may be quite different between the two experiments at the 2-week cold time point (total RNA and chromatin bound RNA were measured in separate experiments, and we have previously observed highly variable changes in *FLC* expression at this time point), this would still not explain the lack of a decreasing trend in the chromatin-bound RNA profile across the intron.

Such a discrepancy between predicted and measured intronic chromatin-bound RNA could, however, result from lariat retention in the chromatin-bound fraction, i.e., some of the intron lariat component also contributing to the chromatin-bound RNA fraction. We therefore modified the model for chromatin-bound intronic RNA to incorporate a partial contribution to this fraction from the intron lariat, consistent with possible intron retention in the chromatin while being degraded. We further assumed that this contribution is unchanged in the cold. With this modified model for chromatin-bound intronic RNA and using the tiling data for the NV and 2W time points, we performed model fitting using our nested sampling approach (see Supplementary Information) and using a modest 0.8-fold reduction in PolII consistent with the termination index at this timepoint (0.75). We obtained good model fits to both total RNA and chromatin-bound RNA profiles (Fig. 4B,C), and inferred the following model parameters (Fig. S8): fold change (FC) in frequency of productive transcription initiation (2W/NV), fold change (FC) in intron processing rate of intron 1 (2W/NV), and fractional contribution to chromatin-bound RNA from the intron lariat. We simultaneously fit the model to the total RNA and chromatin-bound RNA profiles but allow the FC in initiation to vary between the two profiles, to allow for possible differences between separate experiments. The estimated FC in intron processing rate was 6.8±3.4 (mean ± s.d.), indicating a significant increase in intron processing rate at 2W cold. The estimated change in initiation rate varied between the total RNA (0.67±0.06) and chromatin-bound data (1.3±0.3), but this is consistent with variability in *FLC* shutdown at the 2W time point previously observed in our datasets. The fractional contribution from the intron lariat component to the chromatin-bound RNA was estimated to be 0.17±0.1. We assumed the mean estimated fractional contribution of 0.17 for all subsequent fits (Table S1).

We then repeated the intron 1 tiling analysis at the 4 weeks cold time point (4W), using a 0.65-fold experimentally derived reduction in PolII speed. We found that the modified model produces good fits to both chromatin-bound and total RNA profiles (Fig. 4B,C). We repeated the nested sampling approach to infer fold changes in the productive transcription initiation and intron processing rates (Fig. S9). The model predicts a further reduction in initiation at 4W, with estimated fold changes from NV of 0.51±0.03 from total RNA and 0.87±0.11 from chromatin-bound RNA, consistent with more *FLC* copies switching to a PRC2 nucleated, silenced state over time. A further increase in the intron processing rate was also estimated at this time point (FC of 9.4±2.8). We then repeated the same analysis for intron 1 total RNA at the 6W and 8W cold time points. We imposed a further reduction in PolII speed to 0.6 of NV speed, due to the increased fraction of *FLC* gene copies with nucleated H3K27me3 at these timepoints, as well as some H3K27me3 spreading by the 8-week timepoint [20]. We found that the model produces good fits, predicting further reductions in initiation but little further increase in the intron processing rate (Fig. S10). We also repeated the same analysis for intron 1 total RNA at the 7-day post-cold time points after each of the cold treatments. Here, we assumed further reductions in PolII speed at these timepoints, corresponding to increasing fractions of *FLC* copies with spread H3K27me3 [27–29]. We used fold changes (to NV) of 0.6 for 2WT7, 0.5 for 4WT7 (consistent with the termination index fold change at 4WT10 (Fig. 3C)), and 0.3 for 6WT7 and 8WT7. If we assume that the PolII speed in the H3K27me3 spread state is ∼10 fold lower than in the active state [12], and that all *FLC* copies are in one of these two states, the above fold changes correspond to the following fractions of *FLC* copies in the H3K27me3 spread state: ∼0.4 at 2WT7, ∼0.6 at 4WT7, and ∼0.8 at 6WT7 and 8WT7. These fractions were chosen to be consistent with the relative H3K27me3 enrichment over *FLC* measured at these timepoints, where the enrichment was observed to approximately double between 2WT7 and 6WT7/8WT7, and was intermediate at 4WT7 [20]. Significantly, at all of these post-cold time points, the model produced good fits, predicting a recovery in the intron processing rate fold change, i.e., a reduction relative to its cold-enhanced values (Fig. S11), while the productive transcription rate remained lowered as would be expected for epigenetic Polycomb silencing. Thus, the model fits indicate that the enhanced intron processing rate is a cold-specific effect. Overall, these results explain the differences in dynamics of spliced and unspliced *FLC* shown in Fig. 1A, demonstrating that the reason for the larger reduction in unspliced *FLC* in the cold is a cold-specific increase in the intron processing rate.

Previous analysis of short-term cold treatment using plaNETseq has found the opposite phenomenon, with 3 hours of cold resulting in slower splicing kinetics which then recover to pre-cold levels after 12 hours [23]. Since these findings seem to contradict the above results, we assayed the *FLC* spliced to unspliced ratio at higher time resolution (including multiple time points within the first 12-hour phase of the treatment), and extending into long-term cold. We found that indeed this ratio reduces in the first 3 hours, returns to NV levels after around 6 hours, and then shifts to a higher ratio beyond 2 days of cold (Fig. S12). These results are consistent with the transient slowing down of splicing kinetics observed for short-term cold treatment in Kindgren et al. [23]. These results also clearly distinguish two phases of cold treatment in the splicing kinetics: short-term cold stress and long-term vernalization.

## DISCUSSION

Transcriptional regulation can occur at any point in the transcription cycle. While repression at the level of transcription initiation and promoter proximal release is widely reported, the role of other co-transcriptional processes in regulating transcriptional output has received less attention. In this work we report on the transcriptional repression of *FLC* during vernalization taking advantage of mathematical modelling, a vital tool needed to disentangle the underlying transcriptional complexities. Combining experimental data with our model of *FLC* transcription has provided key insights into *FLC* downregulation and quantifies changed dynamics of multiple co-transcriptional processes.

Previous analysis of transcription dynamics at *FLC* in plants grown at warm temperatures has described a transcription-coupled repression mechanism [10, 11] where proximal polyadenylation induces H3K4me1 removal that feeds back to reinforce proximal termination. This graded transcriptional repression determines the timing of the subsequent switch to Polycomb silencing [9, 11], a silenced state which involves co-ordinated changes in transcription initiation and elongation rates [12]. Since cold-induced *FLC* silencing during vernalization is also a Polycomb-mediated mechanism we first thought that cold would promote proximal polyadenylation of potentially both strands at *FLC*, thereby repressing transcription and permitting a transition to cold-induced Polycomb silencing in a similar transcription-coupled repression mechanism. However, our data reveal that transcriptional repression of *FLC* during cold exposure is different from that in the warm, with little or no role for premature termination. This is supported by the lack of vernalization phenotypes of mutants in this pathway such as FCA and FLD [30]. Instead, *FLC* transcriptional repression in the cold can be explained almost entirely by a combination of reduced productive transcription initiation and an enhanced splicing rate.

We have attributed the changes in productive transcription initiation and splicing in the cold to co-transcriptional repression. However, it is also the case that Polycomb silencing is occurring simultaneously. Nevertheless, particularly at earlier time points in the cold (2W) there is only a small contribution from Polycomb silencing. At later time points, Polycomb repression becomes increasingly important. However, as there will be much lower transcription in the Polycomb silenced state, the spliced and unspliced RNAs are likely to be produced predominantly by those *FLC* alleles that have not switched to the Polycomb silenced state. Hence, measurements of spliced to unspliced RNA will reflect the co-transcriptional repression mechanism even at later time points. However, the same will not be true for the PolII speed, for example, which is greatly reduced in the Polycomb silenced state. Hence, an important role for Polycomb in reducing PolII speed has been built into our analysis methodology.

The most counter-intuitive aspect of our findings is that the splicing rate is enhanced by weeks of cold temperature exposure. This contrasts with previous studies concluding that very short-term cold (-3 hours) decreases splicing kinetics [23]. However, these acclimate to NV levels by 12 hours of cold exposure before as we show, increasing after even longer times in the cold, clearly demarcating short-term cold stress from a long-term cold response. Many more studies will be required to analyse splicing dynamics over time in different subsets of genes. However, determination of the rate of splicing is challenging and until recently typically has been assayed through the ratio of unspliced and spliced transcripts. This analysis assumes that there are no differential effects on the turnover dynamics of the two types of transcripts. We demonstrate that this assumption is correct for *FLC* when comparing the warm to relatively long-term cold. Nevertheless, it will be important to return to data collected in other studies to see whether such a constant mRNA half-life is a common feature.

The next problem will be to determine what factors explain the increased splicing rate in long-term cold. Our recent genetic screen identified a role for the spliceosome component SMU1 in co-transcriptional *FLC* repression in the cold [31]. *COOLAIR* proximal intron splicing was promoted by SMU1 in the cold, however *FLC* intron splicing was unaffected. Increased *FLC* splicing could also be connected to changed PolII elongation rate, known to be coupled with splicing [32]. Slow elongation provides a greater opportunity for particular splice sites to be used and has thus been implicated in alternative splicing examples [4, 5]. For *FLC*, however, changes in the transcription elongation rate in the cold appear to be smaller: at 2 weeks cold, for example, PolII speed is reduced by only a factor of ∼0.8, but with at least a 2-fold change in the splicing/intron processing rate. Whether such a small relative change contributes to the enhanced splicing rate at *FLC* in the cold remains to be definitively determined. Further work is also required to analyse changes in PolII CTD phosphorylation at *FLC* in plants exposed to different conditions.

The interconnectivity of different steps in the transcription cycle provides the complexity to differentially regulate transcript abundance of many genes, allowing multiple avenues for finetuning transcriptional output by the environment. This has been central to the regulation of *FLC* and may explain why it is so sensitive to generic transcriptional regulators. Considerable expression variation in *FLC* expression has been selected widely to underpin vernalization, life-history strategy and inflorescence structure [7, 33]. Thus, understanding how natural *cis*-based polymorphisms at *FLC* impact each aspect of the *FLC* transcription cycle will be an important step towards understanding the evolution of genotype-phenotype interactions in changing environments.

## Acknowledgements

We thank Sebastian Marquardt and Peter Kindgren for help in establishing plaNETseq. We also thank all members of the Dean and Howard labs, and Shuqin Chen for excellent technical assistance. This work was funded by European Research Council Advanced Grant (EPISWITCH, 833254), Wellcome Trust (210654/Z/18/Z), and a Royal Society Professorship (RP\R1\180002) to CD, and BBSRC Institute Strategic Programmes (BB/J004588/1 and BB/P013511/1) and EPSRC/BBSRC Physics of Life grant (EP/T00214X/1) to MH and CD. Maria Sindalovskaya was supported by the John Innes Foundation PhD Programme. We thank the JIC Informatics platform for support in analysing the plaNETseq data and the JIC Bioimaging facility for supporting the smFISH experiments.

## Author contributions

Conceptualization, R.M., G.M., M.H., and C.D.; Methodology, G.M., R.M., G.M., S.D., R.B., H.Y., M.H., and C.D.; Software, R.M., M.S., and G.M.; Validation, R.M., R.B., H.Y.; Formal Analysis, G.M. and M.H.; Investigation, G.M.,R.M., S.D., R.B., and H.Y.; Data Curation, G.M., R.M., M.S. and S.D.; Writing – Original Draft, G.M., R.M., C.D., and M.H.; Writing – Review & Editing, G.M., R.M., C.D., and M.H.; Visualization, G.M., R.M., M.S., S.D.; Supervision, C.D. and M.H.; Funding Acquisition, C.D. and M.H. R.M. performed the plaNETseq experiments, *FLC* proximal termination experiments, intron tiling experiments on chromatin bound RNA, and analysed the data. S.D. performed the smFISH experiments to measure mRNA half-life. R.B. performed the high temporal resolution timecourse measurement of *FLC* transcriptional output in the early cold phase. H.Y. performed the *FLC* intron tiling experiments for total RNA. G.M. performed the mathematical modelling, parameter inference, and analysed the data from all experiments. M.S. analysed the plaNETseq data.

## Data availability

Raw read data generated by plaNETseq is available under the SRA Bioproject accession number PRJNA1144665. The data used to generate all plots apart from the plaNETseq plots are available at: https://doi.org/10.5281/zenodo.15477621

These data include an Excel file containing the quantified mRNA levels generated from the smFISH experiment. Supplementary tables containing primer information and smFISH probe sequences are provided as supplementary files and are also available at: https://doi.org/10.5281/zenodo.15477621

Images generated for the smFISH analysis have been uploaded to the EMBL-EBI BioStudies repository and can be accessed at: https://www.ebi.ac.uk/biostudies/studies/S-BSST2030

## Code availability

Python code used to perform parameter inference for the intronic RNA model, to plot *FLC* time course data and mRNA decay data, and all the datasets required for these analyses are available at: https://doi.org/10.5281/zenodo.15477621 The python code is provided in Jupyter notebooks, while the datasets are in the form of text files and Excel files. R code used to analyse and plot the plaNETseq data is also available at https://doi.org/10.5281/zenodo.15477621 in the form of an R markdown file.

## Methods

### Plant materials and growth conditions

All seeds were surface sterilised and sown on Murashige and Skoog (MS) agar plates without glucose. Plates were stratified for 3 days at 4°C. Non-vernalized (NV) plants were grown in long day (LD) conditions (16-h light, 8-h dark with constant 20°C) for 10 days before harvesting. Vernalized plants (WV) were subsequently transferred to short day conditions (8-h light, 16-h dark with constant 5°C) for specified time regiments. For post-cold time points, vernalized plants were transferred to 22°C LD for 7- or 10-days growth (WxT7, WxT10). All plant materials are in a Columbia-0 (Col-0) background with introgression of an active *FRIGIDA* allele from the Sf2 background, described previously [34]. NRPB2-FLAG lines have been previously described and were crossed into the Col*FRI* background containing the active *FRI* allele [23, 35].

### smFISH

Col*FRI* seeds were sown on GM minus glucose media and stratified for 3 days at 5 °C. Non-vernalized (NV) seedlings were grown for 7 days at 22 °C under long day conditions. For 2-week vernalized (2W) plants, seedlings were pre-grown under the same conditions for 7 days, then transferred to 5 °C under short day conditions for 14 days. Whole seedlings were either transferred to Actinomycin D (final concentration 20 µg/ml; Invitrogen, Cat: A7592) or DMSO control plates and returned to their respective temperature conditions. *FLC* mRNA was detected after 0, 2, 4, and 6 hours by smFISH, as described in [36]. Briefly, seedlings were fixed for 30 mins in 4% methanol-free formaldehyde (Sigma-Aldrich, Cat: P6148), then washed three times with 1x PBS. Root tips were protected during manual squashing and the slides were immersed in liquid nitrogen for ∼10 secs. After coverslip removal, samples were dried for 1.5 hr then immersed in 70% EtOH for 1 hr. Samples were equilibrated in 200 µL Stellaris Wash Buffer A (LGC Biosearch, Cat: SMF-WA1-60) containing 10% deionized formamide (Thermo Fisher, Cat: AM9344). Probe hybridization was performed with 150 µL Hybridization Buffer (LGC Biosearch, Cat: SMF-HB1-10) containing 10% formamide and *FLC* mRNA probes diluted to a final concentration of 250 nM. Slides were incubated overnight at 37 °C in the dark. Next day, unbound probes were removed and 150 µL Wash Buffer A was incubated on each slide for 30 mins at 37 °C followed by a 30 min incubation of 150 µL of 1µg/mL DAPI (Sigma, Cat: MBD0015), diluted in Wash Buffer A. Samples were washed for 5 mins in 200 µL Wash Buffer B (LGC Biosearch, Cat: SMF-WB1-20) and mounted in Vectashield mounting medium (Sigma, Cat: H-1000).

Images were acquired on a Zeiss Elyra PS1 inverted wide-field microscope using a 100x oil immersion objective (NA 1.46) and an Andor iXon 897 camera, controlled by Zen 2.3 (Black) software. Z-stacks with 0.2 µm intervals were collected using 642nm and 405 nm lasers to excite Quasar 670-labeled probes and DAPI, respectively. Emission was collected at 655 -710nm (Quasar 670) and 420 - 480nm (DAPI). Image analysis was performed using FIJI [37], as described in [36]. Briefly, the Freehand Selection Tool was used to define regions of interest (ROIs) on maximum intensity z-projections. The “Measure” function recorded cell area, and the “Find Maxima” command with optimised prominence settings was used to quantify mRNA foci. The foci counts per unit area per cell obtained from this analysis was used to estimate the mRNA half-life. To do this, an exponential decay curve was fit to the counts data at the 2hr, 4hr, and 6hr timepoints, avoiding the T0 timepoint to allow time for ActD penetration into plant tissue. Separate fits were performed for the NV and 2W samples, using an ordinary least squares approach, implemented using the numpy.linalg.lstsq function available in the Python library NumPy (Python code provided – see Code availability). In each case, the log-transformed count data was used to estimate two parameters - a slope and an intercept. The standard error of the least squares estimate was computed using the residual variance and the covariance matrix of the regressors.

### CB-RNA preparation

Chromatin-bound RNA (CB-RNA) was isolated as described (Zhang et al., 2022) [38]. Briefly, 3g of non-crosslinked material was homogenised in Honda buffer (0.44 M sucrose, 1.25% (wt/vol) Ficoll, 2.5% (wt/vol) Dextran, 20 mM HEPES pH 7.4, 10 mM MgCl_2_, 0.5% Triton X-100, 2 mM DTT) supplemented with 20 U/mL RNase inhibitor (RNAseIn; Promega, Cat:N2515) and proteinase inhibitor (cOmplete; Roche, Cat:04693116001) and 100 μg/ml yeast tRNA (Roche, Cat:10109509001), before weighing the pellet and resuspending with equal volume of nuclei resuspension buffer (50% Glycerol, 0.5 mM EDTA, 25 mM Tris–HCl pH 7.5, 100 mM NaCl, 2 mM DTT, 20 U/mL RNase inhibitor, 1xproteinase inhibitor, 100 μg/ml yeast tRNA). The resuspended pellet was diluted first with two volumes wash buffer (25 mM Tris–HCl pH 7.5, 300 mM NaCl, 1 M Urea, 0.5 mM EDTA, 1% Tween 20), pipetting 20 times and incubating on ice for 1 minute. Resuspending was performed again with resuspension buffer and repeating with 1 volume of wash buffer, pipetting 20 times. After spinning down the chromatin (12000g for 2 minutes), CB-RNA was isolated by homogenising in 1ml TRIzol (Invitrogen). After addition of 0.2 volumes chloroform, samples were vortexed and centrifuged at 14000g and the aqueous phase isolated. To purify chromatin-bound RNA from the aqueous phase, 0.8 volume of ethanol was added and RNA was isolated using the Qiagen RNeasy mini kit (74104) following the manufacturers protocol. To remove genomic DNA contamination, isolated RNA was treated with TURBO DNase (Invitrogen, Cat:AM1907) and purified by additional column cleanup using Qiagen RNeasy miniprep kit following the manufacturer’s recommendations.

### Gene expression analysis

Total RNA extraction was carried our using a previously described hot phenol method [39]. Genomic DNA contamination was removed by DNAse treatment (TURBO DNASE, Invitrogen, Cat:AM1907). cDNA was synthesised for both purified total and chromatin-bound RNA using gene specific RT primers (Supplemental Table 2) and Superscript IV reverse transcriptase (Invitrogen, Cat:18090050), followed by qPCR with gene specific primers. For analysis of qPCR, data was normalized to the relevant housekeeping gene.

### plaNETseq

plaNETseq libraries were generated as previously described from NRPB2-FLAG seedlings with and without an active *FRI* allele [23, 35]. In brief, 3g of seedlings were homogenised in extraction buffer (0.4M Sucrose, 10mM Tris-HCl pH 8, 10mM MgCl2, 5mM BME, 1x proteinase inhibitor (cOmplete; Roche, Cat:04693116001), 20U/ml RNase inhibitor (RNaseOUT; Invitrogen, Cat:10777019)), filtered through miracloth and centrifuged at 5,000 g for 20 min at 4 °C. The pellet was washed once with nuclear wash buffer (0.25M sucrose, 10mM Tris-HCl pH 8, 10mM MgCl2, 0.3% Tween, 5 mM BME,1x proteinase inhibitor, 20U/ml RNase inhibitor) and finally resuspended in NUC3 buffer (1.7M Sucrose, 10mM Tris-HCl pH 8, 2mM MgCl2, 0.15% Tween, 5mM BME,1x proteinase inhibitor, 20U/ml RNase inhibitor), carefully layered over fresh NUC3 and centrifuged at 4 °C for 1 hour. To lyse the purified nuclei and solubilise the chromatin, the pellet was resuspended in lysis buffer (0.3 M NaCl, 20mM Tris-HCl pH 7.5, 5mM MgCl2, 5mM BME, 0.5% Tween,1x proteinase inhibitor, 20U/ml RNase inhibitor) supplemented with 60U of DNase (Roche) and incubated for 30 minutes at 4 °C at 2000RPM. Immunoprecipitation of FLAG tagged PolII complexes was carried out by the lysate at 4°C with gentle rotation for 2 hours with dynabeads pre-coupled with anti-FLAG antibody (Sigma-Aldrich F3165). Beads were then gently washed six times with wash buffer (0.3 M NaCl, 20mM Tris-HCl pH 7.5, 5mM MgCl2, 5mM BME, 1x proteinase inhibitor, 20U/ml RNase inhibitor). To elute nascent transcripts, beads were suspended in 1ml TRIzol (Invitrogen), incubated for 5 minutes, followed by column-based RNA isolation following the manufacturer’s instructions (Direct-zol RNA miniprep kit).

Libraries were prepared from 100ng of purified nascent RNA using the NEXTflex small-RNA-seq library kit v3 (PerkinElmer, Cat:5132-06). After 3’adapter ligation, RNA was fragmented by incubation with alkaline solution (100 mM NaCO3 pH 9.2, 2 mM EDTA) and heated to 95 °C for 5 min [40]. Fragmented 3’ ligated RNA was purified (RNA-clean XP beads; Beckman Coulter), PNK treated (NEB) for 20 min at 37 °C and the RT-primer reannealed (8 mM), before reintroduction into the library preparation at the adapter inactivation step. Libraries were quantified using the Qubit dsDNA HS assay (Invitrogen) and pooled. Target enrichment of *FLC* and target genes was carried out by in-solution target capture using a custom bait panel. 4,861 synthetic 80-nt biotinylated RNA probes were synthesized, complementary to 32 padded gene sequences at 2x bp tiling density (±1kb padding) (Supplemental Table 2) (mybaits; ArborBiosciences). After library amplification, a sequencing-ready pool of indexed enriched libraries were sequenced on an Illumina Xten or the NEXTseq 550 system PE150 at the Beijing Genomics Institute (BGI). Raw reads have been deposited on SRA under the reference PRJNA1144665.

Data analysis was carried out as in Kindgren et al., 2020 [23]. Unique Molecular Identifiers (UMIs) were first trimmed from the read and appended to the read name with UMI-tools v1.1.1 [41], followed by adapter and read quality trimming with trimmomatic v0.39 [42]. R2 reads were mapped to the Arabidopsis genome (TAIR10) with a splice-aware aligner STAR version 2.7.10a [43]. PCR duplicates were filtered from the alignment files with UMI-tools, low mapping quality reads were removed (MAPQ>10 samtools v1.9 [44]) and reads were flipped to restore the original RNA read strand orientation. Read 3’ends that overlap with 5’ and 3’ splice sites (and likely represent co-transcriptional splicing intermediates) were removed before generating strand-specific coverage files for visualisation of nascent transcripts.

