## Supplementary Information for "Co-transcriptional splicing changes combine with reduced productive transcription initiation for cold-induced repression of *FLC*"

[1] Department of Cell and Developmental Biology, John Innes Centre, Norwich Research Park, Norwich NR4 7UH, UK
[2] Department of Computational and Systems Biology, John Innes Centre, Norwich Research Park, Norwich NR4 7UH, UK
[3] Present address: State Key Laboratory of Hybrid Rice, College of Life Sciences, Wuhan University, Wuhan, 430072, China
[4] Present address: The New Zealand Institute for Plant and Food Research Limited, Christchurch, 8140, New Zealand
†These authors contributed equally to this work

*Corresponding authors

**Mathematical model for the dynamics of *FLC* spliced and unspliced transcripts**
A simple Ordinary Differential Equation (ODE) model is used to describe the dynamics of spliced and unspliced transcripts. The model assumes that all the unspliced transcripts produced are eventually converted to the spliced form. The rate of production of unspliced transcripts is assumed equal to the productive transcription initiation rate. The model equations are as follows:

$$\frac{d[FLC]_{uns}}{dt} = k_{init} - k_{spl}[FLC]_{uns}$$

$$\frac{d[FLC]_{spl}}{dt} = k_{spl}[FLC]_{uns} - k_d[FLC]_{spl}.$$

Here $[FLC]_{uns}$ and $[FLC]_{spl}$ represent the concentration of unspliced and spliced forms respectively. $k_{init}$ represents the productive transcription initiation rate, $k_d$ the degradation rate constant of the spliced form, and $k_{spl}$ the rate constant of unspliced to spliced conversion, i.e., the splicing rate.

At steady state, the spliced and unspliced levels can be computed as follows:

$$[FLC]_{spl} = \frac{k_{init}}{k_d}$$

$$[FLC]_{uns} = \frac{k_{init}}{k_{spl}}.$$

Hence, the spliced to unspliced ratio is given by:

$$R_{spl/uns} = \frac{[FLC]_{spl}}{[FLC]_{uns}} = \frac{k_{spl}}{k_d}.$$

As the degradation rate does not change between cold and non-vernalized (NV) conditions, the change in this ratio between cold conditions and NV can then be used to calculate the fold change in splicing rate between conditions:

$$FC_{spl} = \frac{R_{spl/uns}^{cold}}{R_{spl/uns}^{NV}} = \frac{k_{spl}^{cold}}{k_{spl}^{NV}}.$$

**Mathematical model of RNA PolII distribution across the locus**

To infer potential changes in PolII speed using the plaNETseq data, we use a simple model, where the *FLC* locus is assumed to be divided into two regions – a gene body region and a termination window at the 3' end (see schematic in Fig. 4C), with PolII speed being significantly slower in the termination window. The model assumes a constant PolII speed within each window and that all Pol II that initiate productive transcription reach the termination window. The total steady-state PolII within each window is then given by:

$$Pol\ II_{body} = \frac{k_{init}}{v_{body}} l_{body}$$
$$Pol\ II_{term} = \frac{k_{init}}{v_{term}} l_{term}.$$

Here $k_{init}$ represents the productive transcription initiation rate, $v_{body}$ and $v_{term}$ represent the PolII speed in the gene body and termination window respectively, and $l_{body}$ and $l_{term}$ represent the length of the gene body and termination window regions respectively. The termination index (as plotted in Fig. 4C) is given by:

$$Termination\ index = \frac{Pol\ II_{term}}{Pol\ II_{body}} = \frac{v_{body} l_{term}}{v_{term} l_{body}}.$$

The fold change in the termination index relative to the non-vernalized timepoint can be computed as:

$$FC_{Termination\ index} = \frac{Termination\ index_{cold}}{Termination\ index_{NV}} = \frac{v_{body}^{cold} v_{term}^{NV}}{v_{body}^{NV} v_{term}^{cold}}.$$

Hence, if $v_{term}$, the PolII speed in the termination window, is assumed to be unchanged between conditions, then the fold change in the termination index is equivalent to the fold change in PolII speed within the gene body region:

$$FC_{Termination\ index} = \frac{v_{body}^{cold}}{v_{body}^{NV}}.$$

**Mathematical model of *FLC* intron 1 processing**

We previously developed a mathematical model to capture variation in steady state RNA levels across an intron, incorporating the effects of productive transcription initiation, PolII speed, intron processing, and intron lariat degradation, and applied it to *FLC* intron 1 [1]. The model also incorporated differences between intronic RNA levels measured in total RNA and chromatin-bound fractions – by assuming that the total RNA fraction included contributions from the intron lariat, in addition to the entire chromatin bound fraction. By combining this model with qPCR measurements of RNA levels tiling across *FLC* intron 1 and examining both chromatin-bound and total RNA fractions, we were able to reveal differences in the initiation rate and PolII speed underlying different transcriptional states of the *FLC* locus. Our analysis revealed that the transcriptional differences between an active state of *FLC* in the mutant *fca-9* and a Polycomb silenced state of *FLC* in the wild-type Col-0 involved a coordinated decrease in both initiation rate and PolII speed.

The model assumes that steady state RNA levels along an intron consist of three contributing fractions: (i) nascent RNA from PolII currently transcribing through the intron, (ii) nascent RNA from PolII that have transcribed past the 3' splice site at the end of the intron, but where the intron lariat has not started being degraded, (iii) intron lariat RNA currently being degraded, see Fig. 4A. The total nascent RNA contribution ((i) and (ii) above) can be computed as:

$$\text{Intron 1 Nascent RNA}(x) = \int_{x}^{\text{IA}} \text{PolII}(y)\, dy + \text{RNA}_{\text{s}}$$

where PolII(y) represents the density of PolII at location y, the variable *x* represents position relative to the *FLC* transcription start site (*x*=0 at the TSS), and IA represents the position of the intron acceptor site relative to the TSS. $RNA_s$ represents the contribution from PolII that have transcribed past the 3' end of the intron ((ii) above).

Assuming a constant PolII speed along the intron and no termination within the intron, PolII(y) will be equal to *F/v*, where *F* represents the productive transcription initiation rate (more specifically, the frequency of PolII transcribing into the intron), and *v* represents the PolII speed across the intron. The contribution $RNA_s$ is equal to $F/k_s$, since these transcripts are being produced at a rate *F* and their contribution to intronic RNA is being removed by the process of splicing followed by lariat degradation. Here the parameter $k_s$ represents the intron processing rate. The spliced intronic RNA is not immediately available for lariat degradation, since it has to first undergo further steps including debranching. Taking into account the time taken for these intermediate steps between intron cleavage and the start of lariat degradation, we note that the intron processing rate $k_s$ in the above model does not just represent the rate of cleavage of the intron at its 3' splice site. Rather $k_s$ in the model is a composite parameter representing the rate of the whole process through which intronic RNA that is initially part of the nascent transcript goes on to become intron lariat RNA subject to degradation. The nascent RNA contribution at a location x within the intron can thus be computed as:

$$\text{Intron 1 Nascent RNA}(x) = F\left(\frac{IA - x}{v} + \frac{1}{k_s}\right)$$

Similarly, the lariat contribution to intronic RNA levels can be computed as follows:

$$\text{Intron 1 Lariat RNA}(x) = F\left(\frac{x - ID}{k_d}\right)$$

assuming a constant rate of lariat degradation in the 5' to 3' direction, with a degradation rate of $k_d$ (bp/s). ID represents the position of the intron donor site relative to the TSS.

The resulting equation for the Wu et al. 2016 [1], model was as follows:

$$\text{Intron 1 RNA}(x)_{\text{Total}} = F\left(\frac{IA - x}{v} + \frac{1}{k_s} + \left(\frac{x - ID}{k_d}\right)\right)$$

for the total RNA fraction.

As discussed above, the parameter $k_s$ is not equivalent to the splicing rate whose fold change was estimated in the main paper using the spliced to unspliced ratio. This measurement was made via an intron-exon junction assay, and so may be directly interpreted as the rate of cleavage at the 3' splice site. However, the two rate measurements – splicing rate and intron processing rate both give consistent increases in the cold versus NV, albeit for different introns, thereby giving confidence that splicing rates are generally increased at *FLC* in the cold.

It is also possible to include 3'-5' lariat degradation in the above model. However, in the current study, introducing 3'-5' lariat degradation at a similar rate to the 5'-3' degradation was found to make the model fit badly. Even in the best fit model, it resulted in a pronounced reduction in the predicted profile from the middle of the intron towards the 3' end. Since the data does not exhibit such a trend, we assume that any 3'-5' lariat degradation must be happening at a much lower rate than 5'-3' degradation. We previously estimated lariat degradation rate using an smFISH based quantitative approach in Ietswaart et al., 2017 [2], where we allowed for degradation in both directions, and the estimated rates were comparable for 5'-3' and 3'-5' degradation. However, in Wu et al. 2016 [1], we found that 3'-5' degradation could be comparable but had to be slower than 5'-3' degradation to allow good model fits to the intron tiling fold change profiles between high transcriptional and silenced states of *FLC*. The reason the model fits in Wu et al. 2016 [1] can accommodate comparable 3'-5' degradation is that there are large fold changes in initiation (~25 fold) and PolII speed (~10 fold) between active and silenced states. In the current study, PolII speed changes (up to ~1.5 fold) and initiation rate changes (up to ~2 fold) between warm and 2W, 4W cold. This makes the model predictions more sensitive to the presence of significant 3'-5' lariat degradation. We therefore neglect it in the model.

The Wu et al. 2016 model assumed that the intron lariat RNA was completely released from the chromatin, i.e., lariat RNA did not contribute to the chromatin-bound RNA fraction in this model. The equation for the intron 1 chromatin-bound RNA was therefore as follows:

$$\text{Intron 1 RNA}(x)_{Chr} = F\left(\frac{IA - x}{v} + \frac{1}{k_s}\right).$$

For this equation to be correct, the intron needs to be cleaved at the 3' splice site before the RNA is released from the locus. According to previous estimates of PolII transcription speed across *FLC* [1, 2], and the subsequent rate of transcript release from the locus, the time available between transcription of the intronic RNA and RNA release from the locus is on the order of 30 min. Indeed, the reported timescales for cleavage of introns is of the order of tens of seconds across different organisms [3], and earlier estimates for *FLC* are around 5 min [2].

We first used the same model and the same methodology as Wu et al. 2016 [1], using intron 1 to examine cold-induced transcriptional repression at *FLC*. However, as explained in the Results section, this model could not capture the experimentally observed trends for RNA levels across intron 1 in both chromatin-bound and total RNA fractions. Specifically, the experimentally measured fold-change profiles (relative to NV samples) in both fractions exhibit an increasing trend from the 5' to the 3' end of the intron. The Wu et al., 2016 model could not capture these trends through a coordinated reduction only in initiation rate and PolII speed (Fig. S4). However, through an increased intron processing rate in the cold combined with reduced initiation rate, the model could capture the 5' to 3' trend  trend for the total RNA fraction. The model was still unable to capture the trend for the chromatin-bound RNA fractions. Importantly, this trend in the experimental data is consistent across cold time-points, and for both total RNA and chromatin-bound RNA fractions.

We therefore developed a modified model by introducing an additional component into the Wu et al. 2016 model – the possibility of partial lariat retention and degradation in the chromatin-bound RNA fraction. Thus, in the modified model, the equation for the total RNA fraction remains the same, while the equation for the chromatin-bound RNA fraction is as follows:

$$\text{Intron 1 RNA}(x)_{\text{Chr}} = F\left(\frac{IA - x}{v} + \frac{1}{k_s} + \text{f}_{\text{lariat}}\left(\frac{x - ID}{k_d}\right)\right),$$

where $f_{\text{lariat}}$ is a parameter constrained to be between 0 and 1, representing the partial contribution from the intron lariat component to the chromatin-bound RNA fraction.

**Model Fitting and Parameter Inference**

Similar to Wu et al. 2016, where we performed quantitative fitting of the model to fold changes in RNA levels between mutants and a wild-type, here we perform quantitative fits to fold changes between time points, specifically between each cold time point and the non-vernalized (NV) time point.

For fitting the intron 1 fold change profiles to both chromatin-bound and total RNA fraction datasets, we fix the basal absolute values of all model parameters corresponding to the non-vernalized (NV) time point and use parameter inference to estimate the fold changes of specific parameters at the cold-treated time points. Thus, the model used for parameter inference is as follows:

$$\text{Fold}(x)_{\text{Total}} = \frac{\text{fc}_{\text{init}}(\text{t})F_{\text{NV}}\left(\frac{IA-x}{v_{\text{cold}}} + \frac{1}{\text{fc}_{\text{proc}}(\text{t})k_s(\text{NV})} + \left(\frac{x-ID}{k_d}\right)\right)}{F_{\text{NV}}\left(\frac{IA-x}{v_{\text{NV}}} + \frac{1}{k_s(\text{NV})} + \left(\frac{x-ID}{k_d}\right)\right)}$$

for the total RNA fraction, and

$$\text{Fold}(x)_{\text{Chr}} = \frac{\text{fc}_{\text{init}}(\text{t})F_{\text{NV}}\left(\frac{IA-x}{v_{\text{cold}}} + \frac{1}{\text{fc}_{\text{proc}}(\text{t})k_s(\text{NV})} + f_{\text{lariat}}\left(\frac{x-ID}{k_d}\right)\right)}{F_{\text{NV}}\left(\frac{IA-x}{v_{\text{NV}}} + \frac{1}{k_s(\text{NV})} + f_{\text{lariat}}\left(\frac{x-ID}{k_d}\right)\right)}$$

for the chromatin-bound RNA fraction. Here, the parameters $\text{fc}_{\text{init}}$ and $\text{fc}_{\text{proc}}$ represent the fold changes in the productive initiation rate and intron processing rate at a given time point relative to the NV values of these rates, see Fig. 4B,C.

Throughout this analysis productive transcription initiation appears only as a fold change between experimental conditions – we therefore do not assume an absolute value for this parameter. PolII speed at *FLC* corresponding to non-vernalized Col*FRI* is assumed to be $v$= 40 bp/s, the estimated mean value for a high transcriptional state of *FLC* from Wu et al. 2016 [1]. Using the median PolII speed estimated in Ietswaart et al., 2017 (11 bp/s), the model could not qualitatively fit the trends in our data. The basal intron processing rate is fixed at 0.002 $s^{-1}$, consistent with the estimate for this parameter from Ietswaart et al., 2017 [2]. Note that, as discussed above, this parameter represents a composite of multiple steps in the model, and therefore should not be treated as equivalent to the rate of intron cleavage at the 3' splice site. The basal 5'-3' degradation rate is assumed to be 3 bp/s, the estimated median value of this parameter from Ietswaart et al., 2017. The inferred parameters for the modified model were the fold changes in initiation and intron processing rates

($fc_{init}$ and $fc_{proc}$), with a fold change reduction in PolII speed that changes over the cold timepoints as described in the text. This is consistent with a cold-induced slowdown suggested by our plaNETseq data (termination index in Fig. 3C), previous quantitative analysis of co-transcriptionally added H3K36me3 [4], and the estimated difference in PolII speed between an active transcriptional state and a Polycomb silenced (H3K27me3 spread) state [1]. The $f_{lariat}$ parameter, representing the lariat component contribution to the chromatin-bound fraction, was inferred using data for the 2 week time point, and held fixed at the estimated value (0.17) for all other time points. Allowing the lariat degradation rate to change in the cold (along with the above inferred fold changes), and inferring this change as an additional parameter, predicted an increase in the lariat degradation rate in the cold, however without significantly improving the model fits (fitted models compared using the Bayesian evidence – see below). We therefore keep the lariat degradation rate fixed at its basal value for all time points. Parameter values used for the *FLC* intron 1 processing model are also shown in Table S1.

We used a Bayesian approach for model parameter inference [5], using a nested sampling algorithm [6] implemented in the Python package Nestle [7]. We used uniform prior distributions for all the inferred parameters, with bounds as specified in the figure captions (Figs. S5-S9). The nested sampling algorithm enables parameter inference by simultaneously computing the posterior distribution of parameters and the Bayesian evidence for the model [8]. The algorithm starts with an ensemble of points in parameter space drawn using the prior distributions, computes the likelihood for each of these samples and iteratively replaces points with the lowest likelihood with new higher likelihood points. At each iteration, the algorithm computes an estimate of the Bayesian evidence, i.e., the integral of the likelihood over the prior distributions, eventually converging to an estimate of the evidence. The termination criterion used by Nestle is based on an estimate of the maximum remaining evidence – the algorithm terminates when the relative contribution from the remaining evidence to the current estimate of the total evidence falls below a threshold [7]. We used the default value of 0.5 for this threshold.

The nested sampling algorithm can be implemented using a variety of methods which differ in how they choose samples from the parameter space [8]. We used the single ellipsoid method [9], which is one of the methods implemented in the Nestle package. To specify the likelihood function, we assumed an additive, Gaussian distributed error in the data (fold change profiles relative to NV time point), with zero mean and standard deviations corresponding to each primer estimated from the three replicates.

1. Wu, Z., et al., *Quantitative regulation of FLC via coordinated transcriptional initiation and elongation.* Proc Natl Acad Sci U S A, 2016. **113**(1): p. 218-23.
2. Ietswaart, R., et al., *Cell-Size-Dependent Transcription of FLC and Its Antisense Long Non-coding RNA COOLAIR Explain Cell-to-Cell Expression Variation.* Cell Syst, 2017. **4**(6): p. 622-635 e9.
3. Alpert, T., L. Herzel, and K.M. Neugebauer, *Perfect timing: splicing and transcription rates in living cells.* Wiley Interdiscip Rev RNA, 2017. **8**(2).
4. Nielsen, M., et al., *COOLAIR and PRC2 function in parallel to silence FLC during vernalization.* Proc Natl Acad Sci U S A, 2024. **121**(4): p. e2311474121.

5. Pullen, N. and R.J. Morris, *Bayesian model comparison and parameter inference in systems biology using nested sampling.* PLoS One, 2014. **9**(2): p. e88419.
6. Skilling, J., *Nested sampling.* Bayesian inference and maximum entropy methods in science and engineering, 2004. **735**: p. 395-405.
7. Barbary, K., *nestle: Pure Python, MIT-licensed implementation of nested sampling algorithms for evaluating Bayesian evidence.* 2014.
8. Ashton, G., et al., *Nested sampling for physical scientists.* Nature Reviews Methods Primers, 2022. **2**(1): p. 39.
9. Mukherjee, P., D. Parkinson, and A.R. Liddle, *A Nested Sampling Algorithm for Cosmological Model Selection*. The Astrophysical Journal, 2006. **638**(2): p. L51.

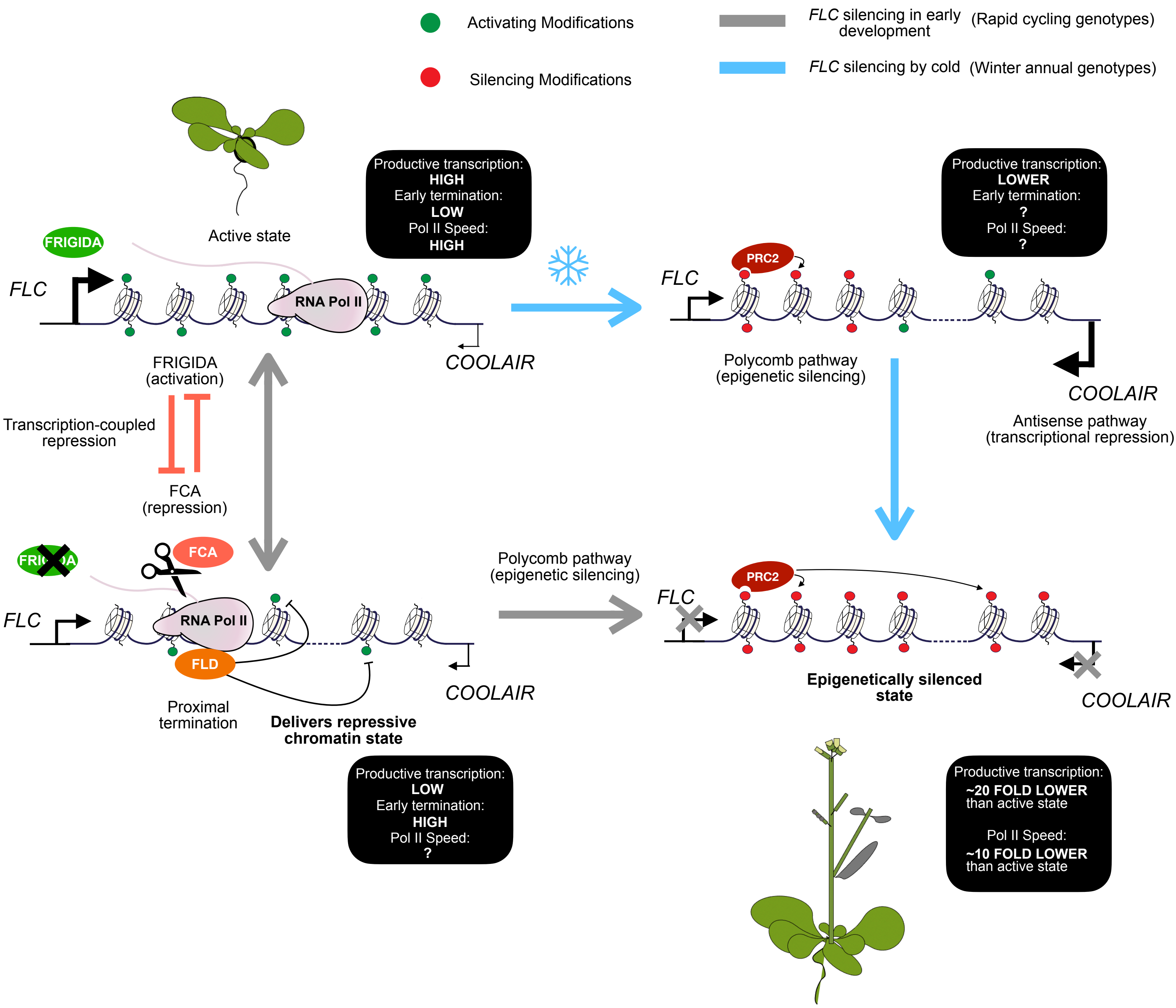


**Figure S1: Schematic summary of transitions in *FLC* transcriptional state and the molecular mechanisms regulating these transitions.**

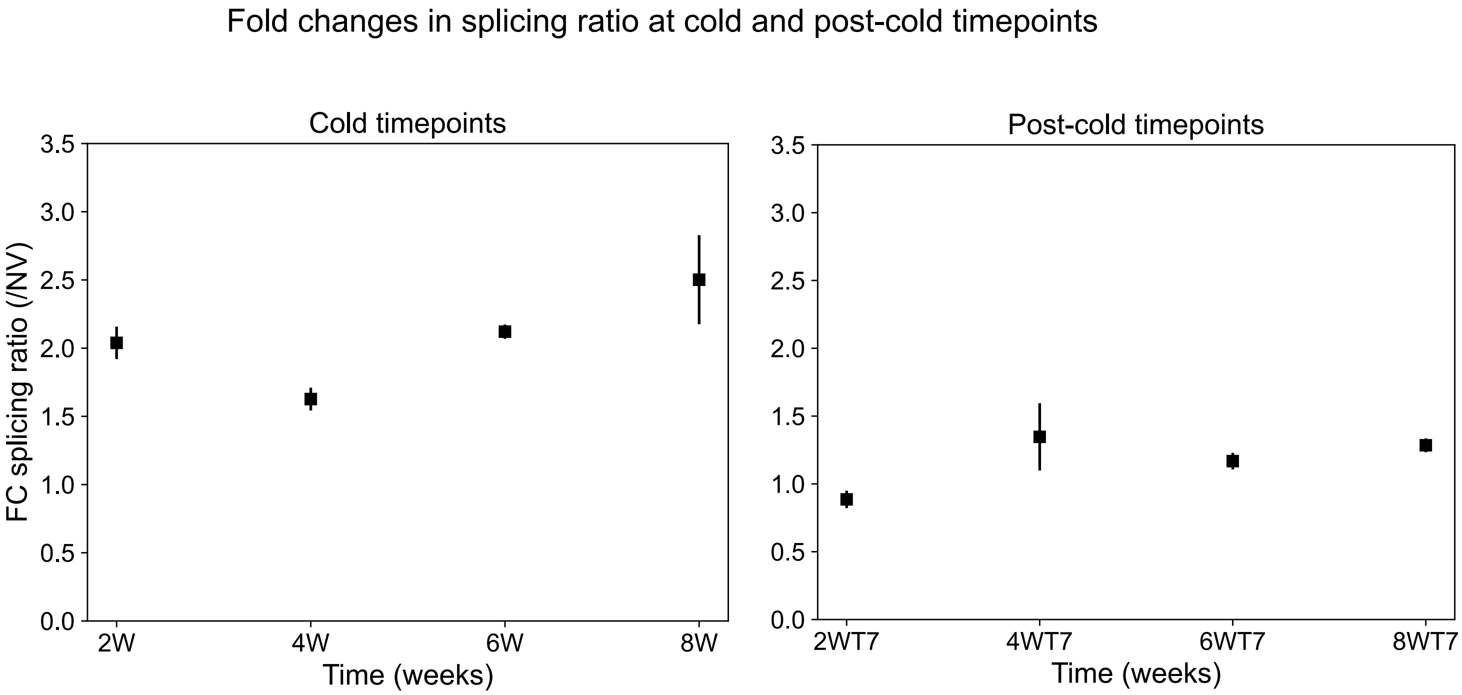


**Figure S2: Fold changes in splicing ratio calculated for all cold and post-cold timepoints.** The fold changes were calculated using the data shown in Figure 1A. See model description for the ODE model showing how this fold change can represent the fold change in splicing rate at these timepoints, relative to the non-vernalised condition (NV). Error bars represent SEM (n=3 biological replicates).

Correlation analysis of plaNETseq samples

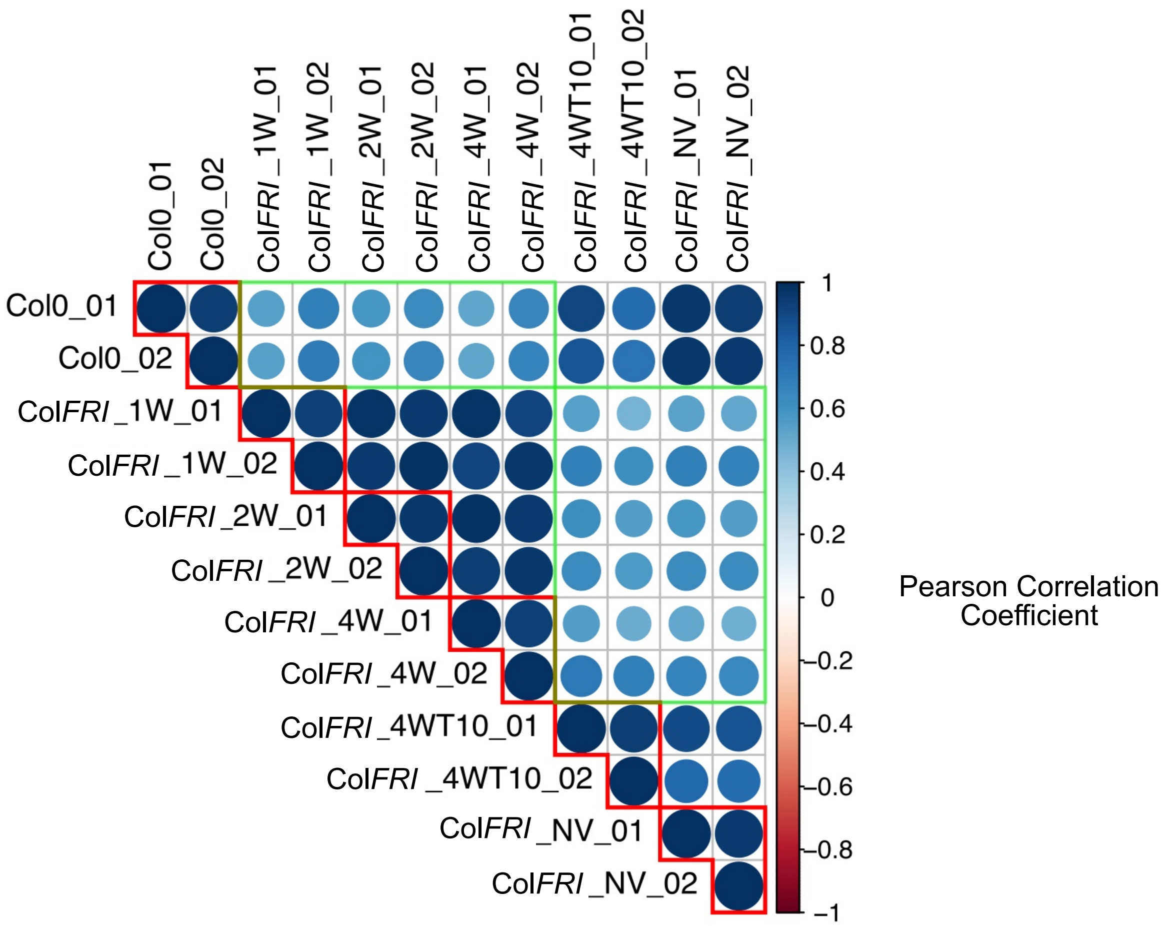


**Figure S3: Correlation analysis of plaNETseq reads after in solution target capture.** Colour intensity and the size of the circle are proportional to the Pearson correlation coefficients between each pair of samples. Red boxes indicate replicate comparisons and green boxes highlight NV and Cold library comparisons. Library correlation remains high for replicates and is reduced when comparing treatment groups.

**Normalized plaNETseq counts of control genes**

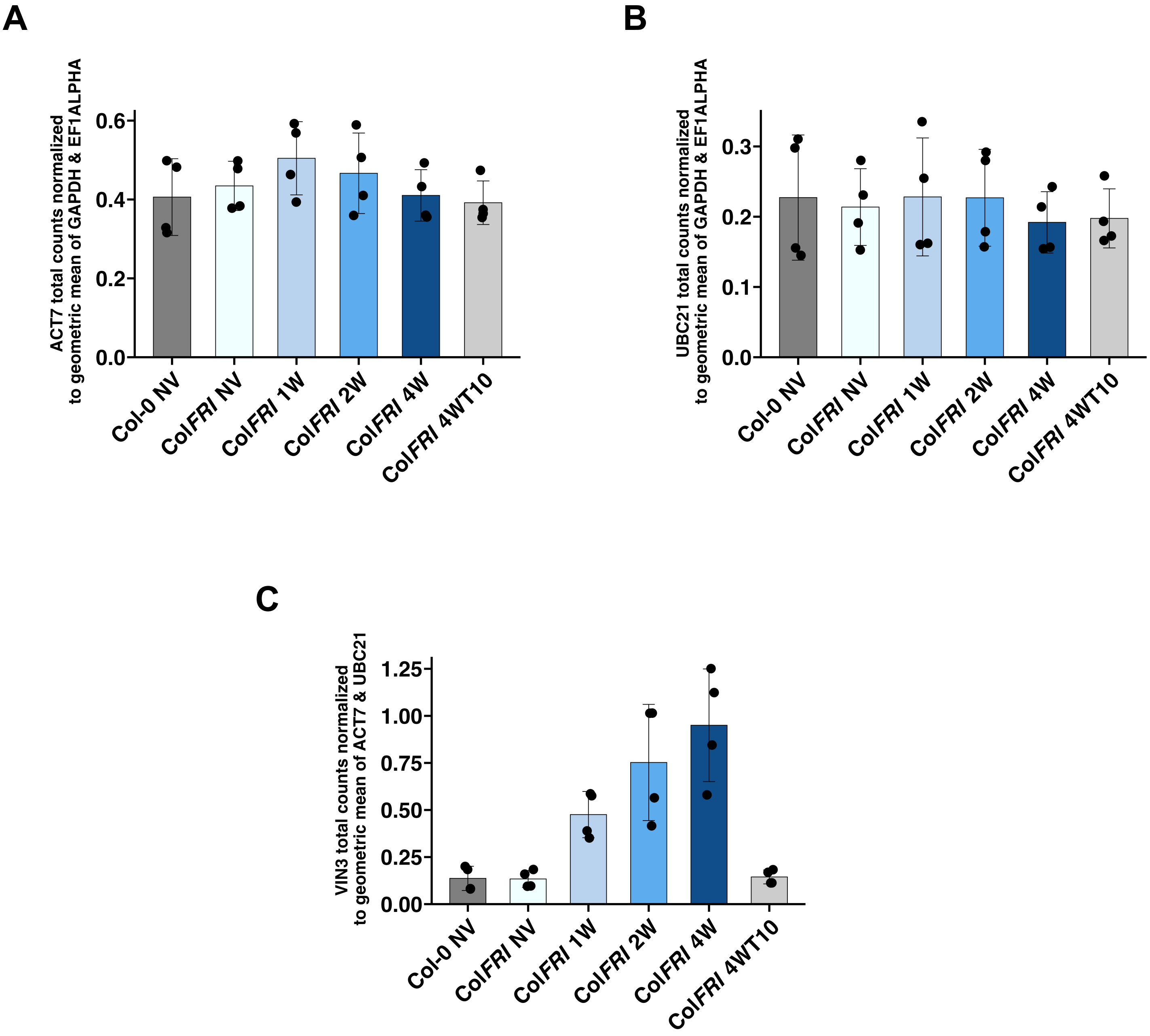


**Figure S4: plaNETseq levels of control genes.** Total plaNET-seq counts of bait-enriched control genes. Mean counts of 4 biological replicates of *ACT7* (A) and *UBC21* (B), normalized to the geometric mean of *GAPDH* and *EF1ALPHA*, and *VIN3* (C), normalized to the geometric mean of *ACT7* and *UBC21*.

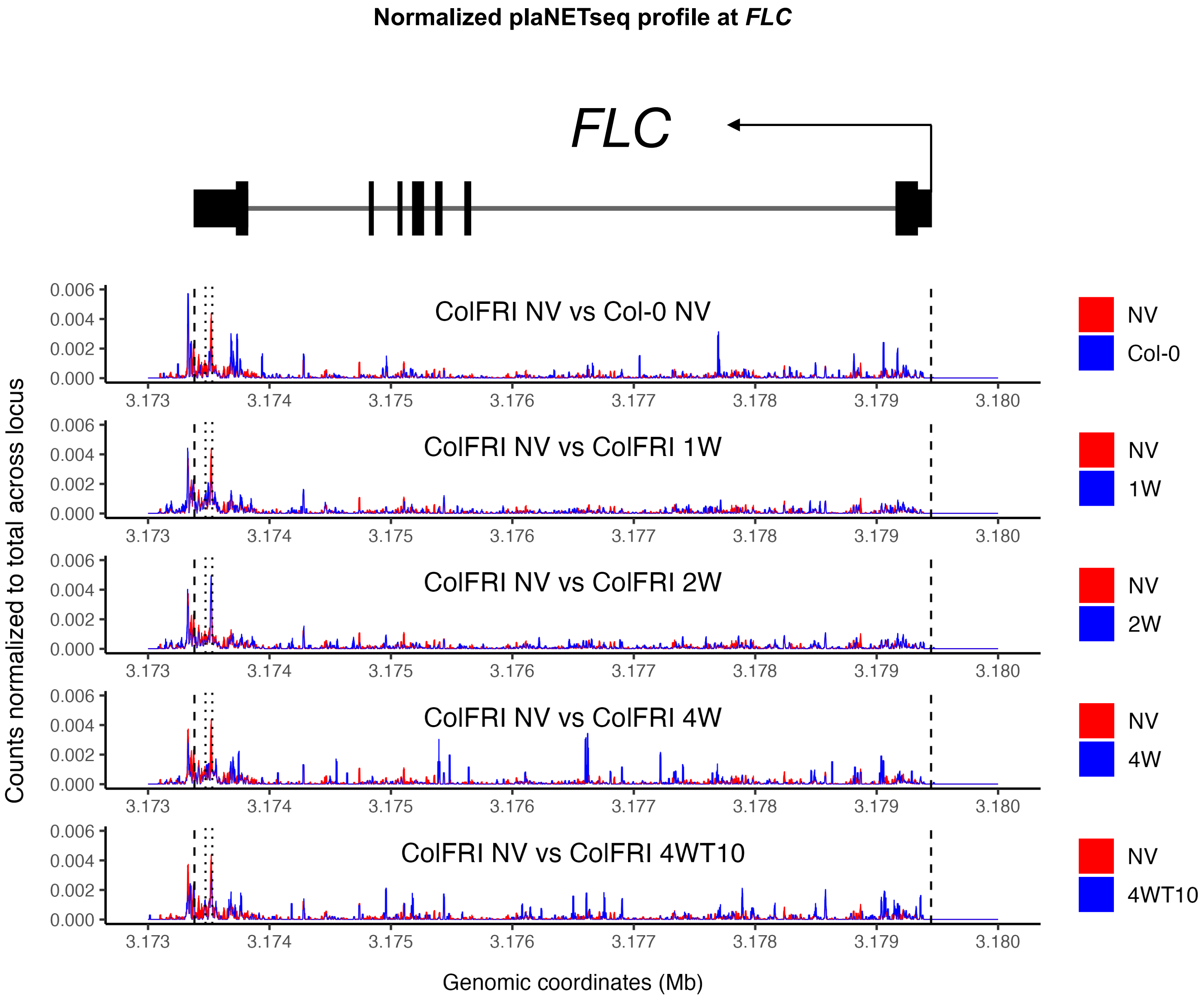


**Figure S5. plaNETseq analysis of the *FLC* locus over a cold time course.** Normalized PolII distributions across *FLC* over the vernalization time series. Mean counts of 4 biological replicates was normalized to the total PolII levels across the locus. Each panel shows a comparison between a cold timepoint and NV conditions in Col*FRI* or between Col-0 NV and Col*FRI* NV. Distributions are plotted as curves and the space between the curves is shaded blue where the cold timepoint or Col-0 NV shows higher normalized PolII level and red where the Col*FRI* NV timepoint shows higher normalized PolII level. *FLC* TSS and the end of 3' UTR (Araport 11 annotation) are represented by black dashed lines and major poly-A sites are shown by black dotted lines. (B) Total plaNETseq counts over *FLC* over the cold timecourse.

Model predictions compared to experimental data (2W/NV) with only productive initiation and Pol II speed allowed to change in the cold

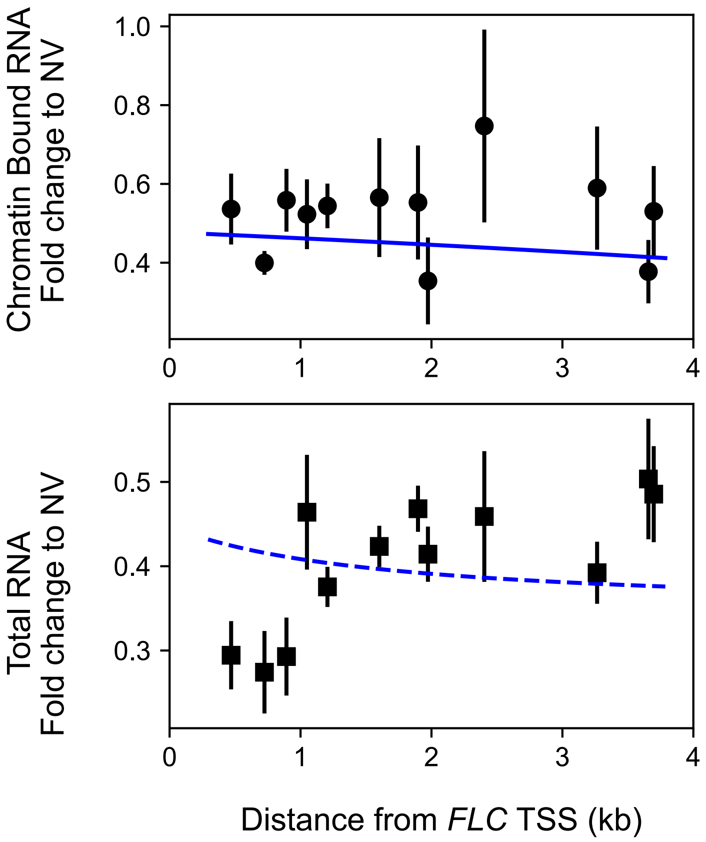


$$FC_{init\ total} = 0.38 \quad FC_{init\ chr} = 0.41 \quad FC_{speed} = 0.5$$

**Figure S6: Model fails to fit measured profiles with only initiation and Pol II speed changes.** Model predictions indicated by blue lines – chromatin bound RNA by solid line and total RNA by dashed line. Experimental datapoints indicated by black dots (chromatin bound RNA) and black squares (total RNA). With only coordinated reduction in initiation and PolII speed, the model fails to even qualitatively capture the trend in the total RNA profile over intron 1 at 2W relative to NV. Best fit shown, with parameter values for fold changes in productive initiation and PolII speed shown. Error bars represent SEM (n=3 biological replicates).

Parameter inference for Wu et al., 2016 *FLC* Intron 1 model using only Total RNA Intron 1 tiling data (2W relative to NV)

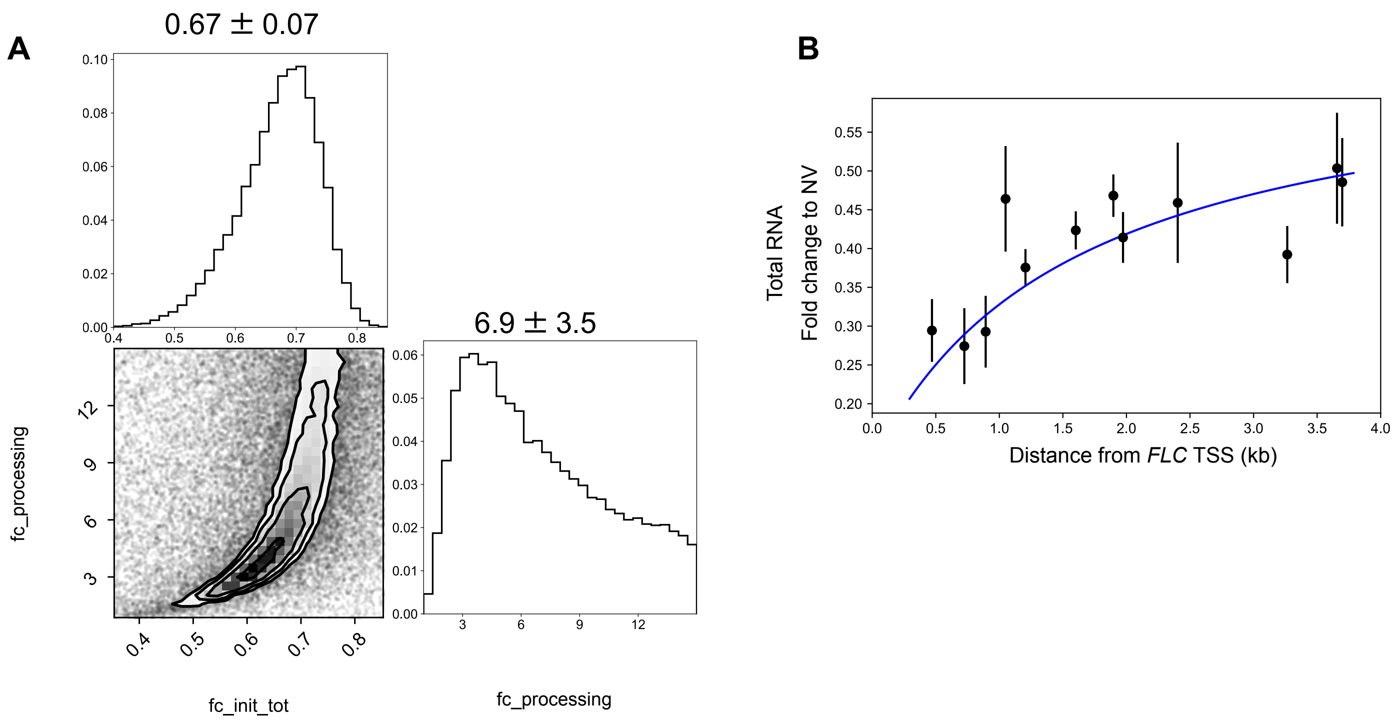


**Figure S7. Estimated posterior distribution of the inferred parameters, obtained using a nested sampling approach, from 2W timepoint Total RNA data.** (A) Two model parameters were inferred using data for the 2W time point. Total RNA data was used to simultaneously constrain the model, and PolII speed was assumed to change by a factor of 0.8. Individual contour plots show two dimensional projections of the final set of parameter samples generated by the nested sampling algorithm. The histograms show the posterior probability of the corresponding parameter values . Mean ± standard deviation of the posterior samples is shown for each parameter. The parameter names are as follows: fc_init_tot denote fold changes in initiation (2W/NV) estimated from the total RNA profile; fc_processing denotes the fold change in intron processing rate (2W/NV); Uniform priors were assumed for all four parameters, within the following bounds: 0<fc_init_tot<2; 0<fc_processing<15; 0<f_lariat<0.4. (B) Model fits using the mean values of the posterior sample distributions for all four parameters.

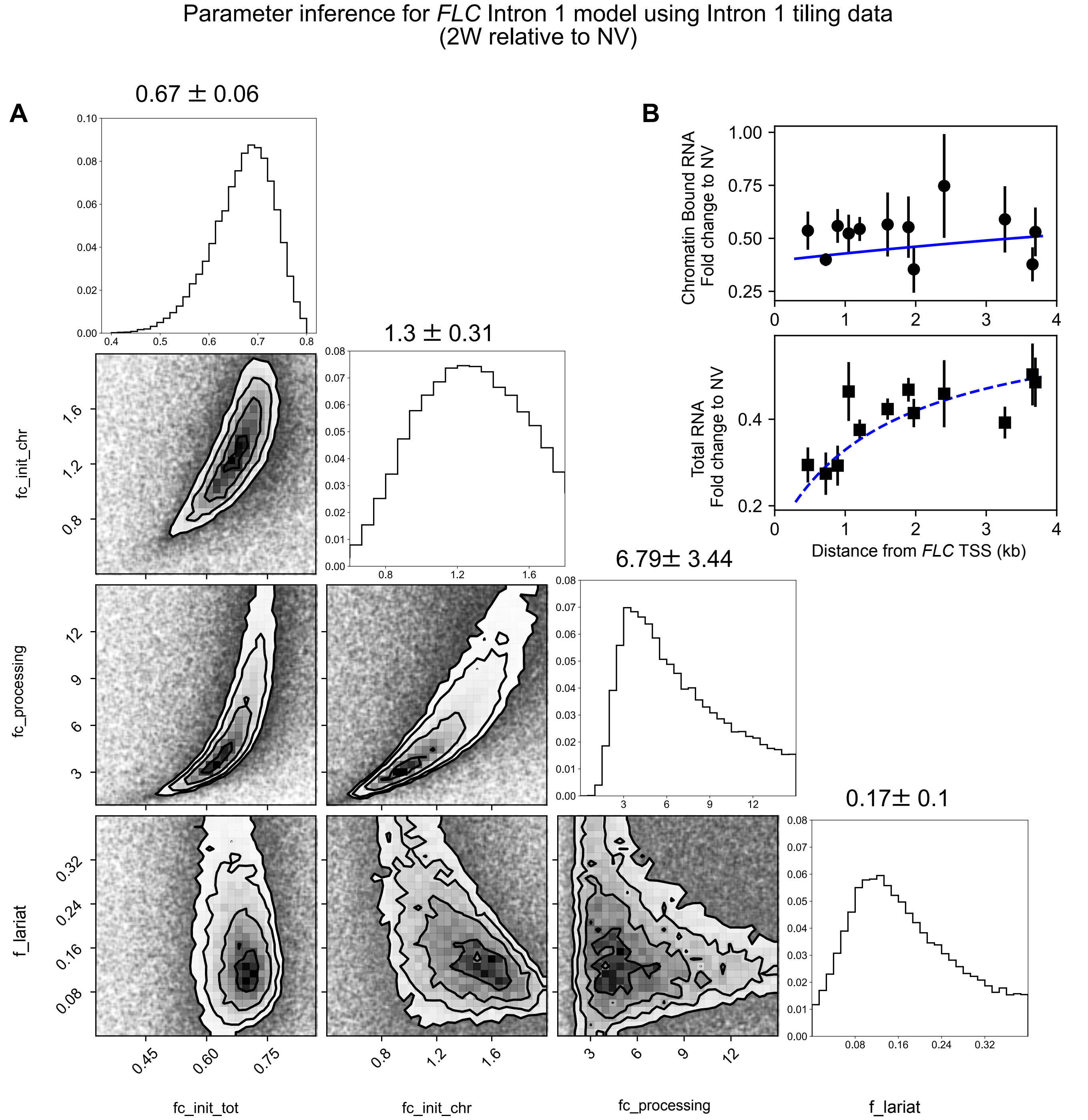


**Figure S8. Estimated posterior distribution of the inferred parameters, obtained using a nested sampling approach, from 2W timepoint data.** (A) Four model parameters were inferred using data for the 2W time point. Chromatin bound RNA data and total RNA data were used to simultaneously constrain the model, and PolII speed was assumed to change by a factor of 0.8 relative to NV. Individual contour plots show two dimensional projections of the final set of parameter samples generated by the nested sampling algorithm. The histograms show the posterior probability distribution of each inferred parameter. Mean ± standard deviation of the estimated posterior distribution is shown for each parameter. The parameter names are as follows: fc_init_tot and fc_init_chr denote fold changes in initiation (2W/NV) estimated from the total RNA and chromatin bound RNA profiles, respectively; fc_processing denotes the fold change in intron processing rate (2W/NV); f_lariat denotes the fractional component of lariat contributing to the chromatin-bound RNA fraction. Uniform priors were assumed for all four parameters, within the following bounds: 0<fc_init_tot<2; 0<fc_init_chr<2; 0<fc_processing<15; 0<f_lariat<0.4. (B) Model fits using the mean values of the posterior sample distributions  for all four parameters (reproduced from Fig. 5).

Parameter inference for *FLC* Intron 1 model using Intron 1 tiling data
(4W relative to NV)

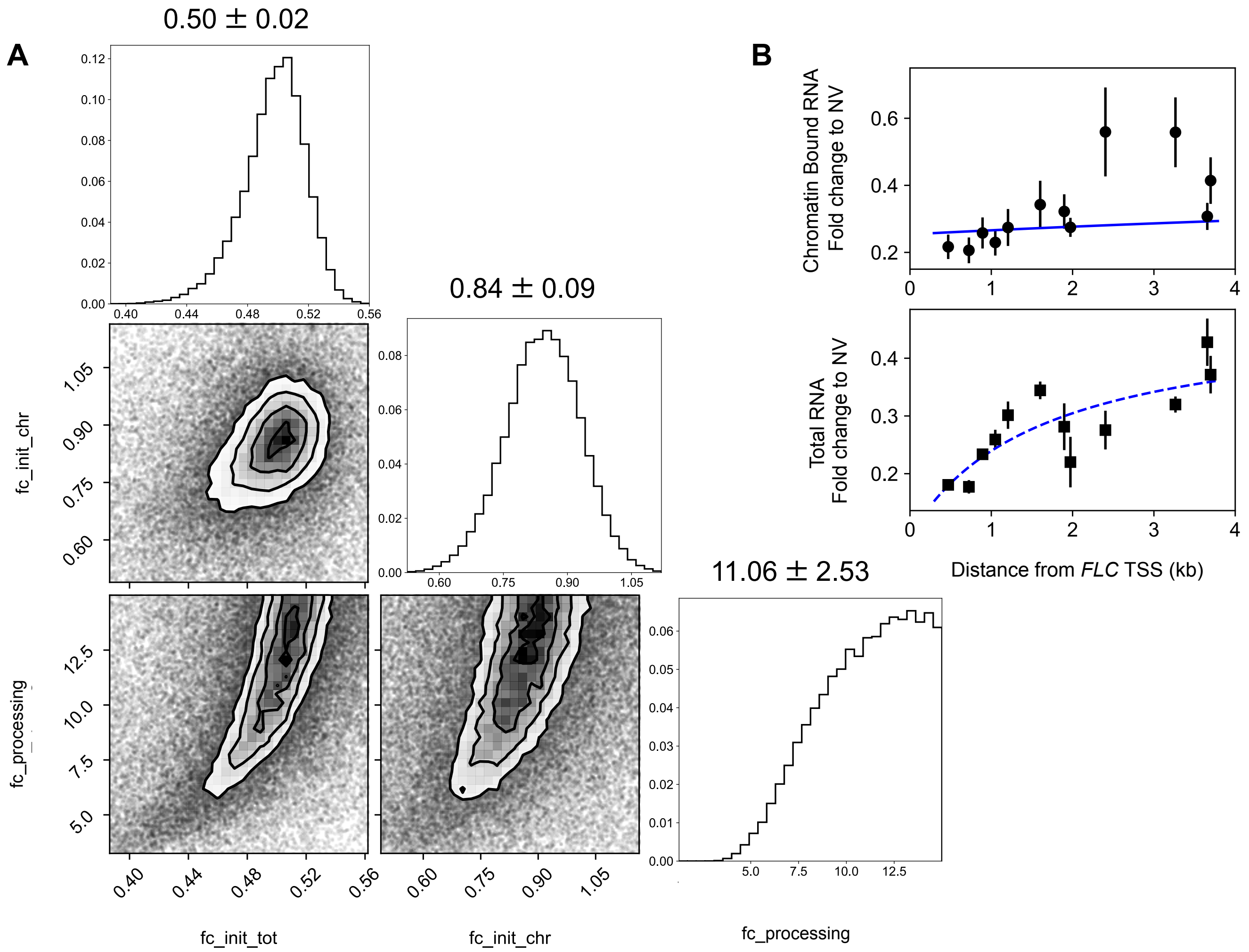


**Figure S9. Estimated posterior distribution of the inferred parameters, obtained using a nested sampling approach, from 4W timepoint data.** (A) Three model parameters were inferred using data for the 4W time point. Chromatin-bound RNA data and total RNA data were used to simultaneously constrain the model and PolII speed was assumed to change by a factor of 0.65 relative to NV. Individual contour plots show two dimensional projections of the final set of parameter samples generated by the nested sampling algorithm. The histograms show the posterior probability distribution of each inferred parameter. Mean ± standard deviation of the estimated posterior distribution is shown for each parameter. The parameter names are as follows: fc_init_tot and fc_init_chr denote fold changes in initiation (4W/NV) estimated from the total RNA and chromatin bound RNA profiles, respectively; fc_processing denotes the fold change in intron processing rate (4W/NV); f_lariat was fixed at 0.17 (the mean value of the posterior distribution estimated from the 2W data). Uniform priors were assumed for all three parameters, within the following bounds: 0<fc_init_tot<2; 0<fc_init_chr<2; 0<fc_processing<15. (B) Model fits using the mean values of the posterior sample distributions for all three parameters (reproduced from Fig. 5).

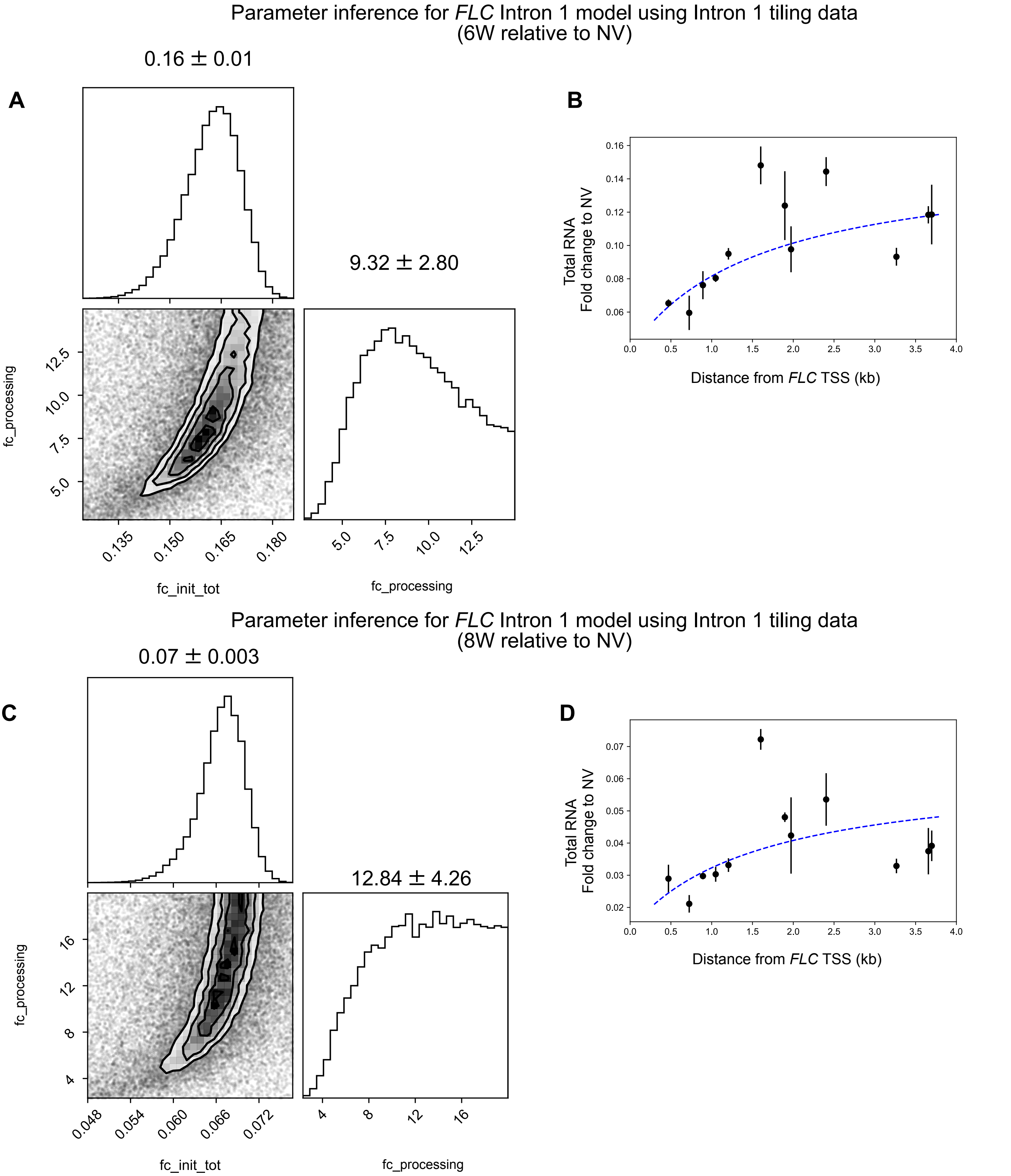


**Figure S10. Estimated posterior distribution of the inferred parameters, obtained using a nested sampling approach, from 6W and 8W timepoint data.** (A,C) Two model parameters were inferred using data for the 6W and 8W time points. Total RNA data was used to constrain the model, and PolII speed was assumed to change by a factor of 0.6 relative to NV. Individual contour plots show two dimensional projections of the final set of parameter samples generated by the nested sampling algorithm. The histograms show the posterior probability distribution of each inferred parameter. Mean ± standard deviation of the estimated posterior distribution is shown for each parameter. The parameter names are as follows: fc_init_tot denotes fold changes in initiation (relative to NV), estimated from the total RNA profile; fc_processing denotes the fold change in intron processing rate (relative to NV); f_lariat was fixed at 0.17 (the mean value of the posterior distribution estimated from the 2W data). Uniform priors were assumed for both parameters, within the following bounds: 0<fc_init_tot<2; 0<fc_processing<15. (B,D) Model fits using the mean values of the posterior sample distributions for the two parameters.

Parameter inference for *FLC* Intron 1 model using Intron 1 tiling data
(Post-cold relative to NV)

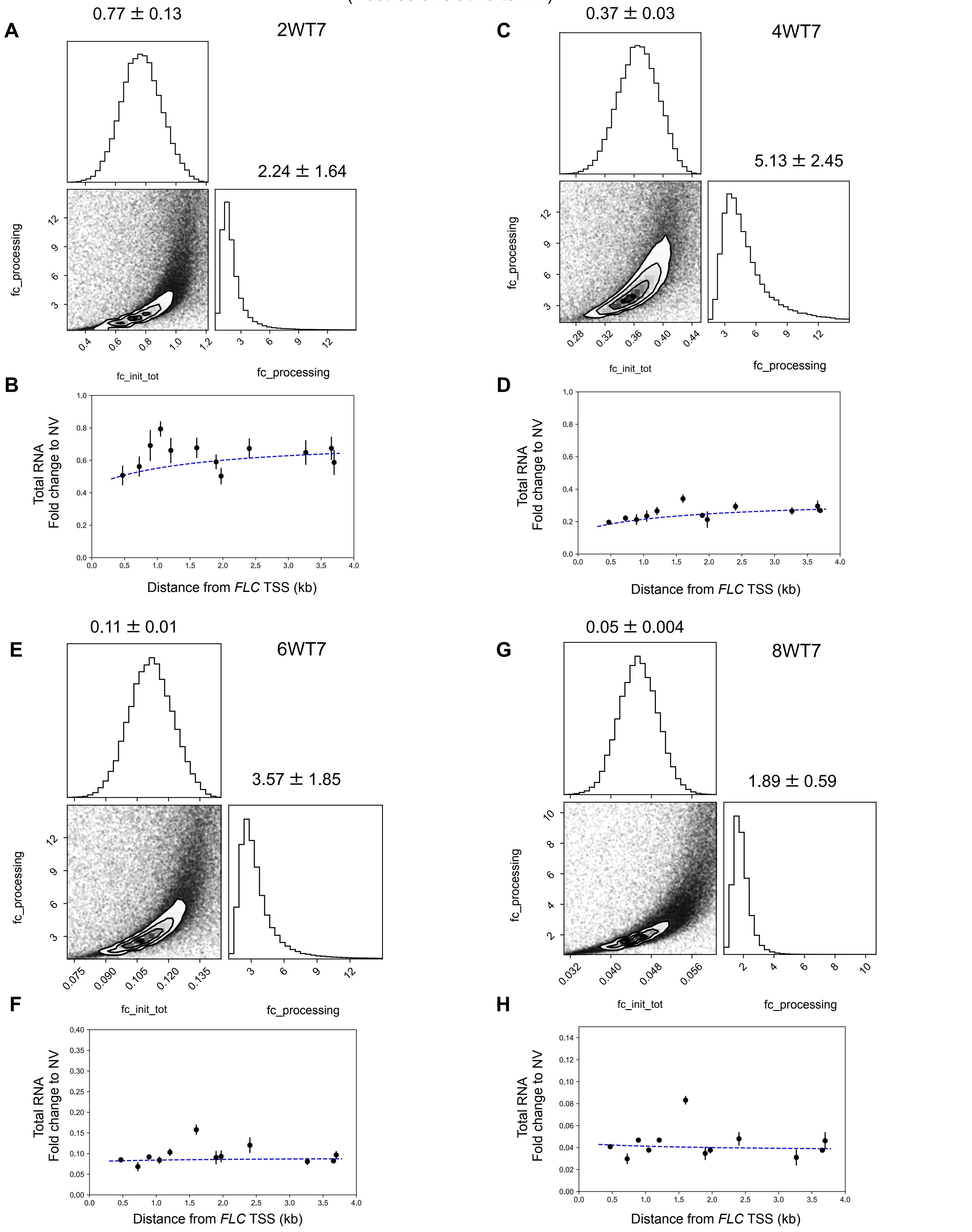


**Figure S11. Estimated posterior distribution of the inferred parameters, obtained using a nested sampling approach, from data for all post-cold timepoints.** (A,C,E,G) Two model parameters were inferred using data for all post-cold time points: 2WT7, 4WT7, 6WT7, 8WT7. Total RNA data was used for parameter inference, and PolII speed is assumed to change by the following factor relative to NV: 0.6 at 2WT7, 0.5 at 4WT7, 0.3 at 6WT7 and 8WT7. Individual contour plots show two dimensional projections of the final set of parameter samples generated by the nested sampling algorithm. The histograms show the posterior probability distribution of each inferred parameter. Mean ± standard deviation of the estimated posterior distribution is shown for each parameter. The parameter names are as follows: fc_init_tot denotes fold changes in initiation (relative to NV), estimated from the total RNA profile; fc_processing denotes the fold change in intron processing rate (relative to NV); f_lariat was fixed at 0.17 (the mean value of the posterior distribution estimated from the 2W data). Uniform priors were assumed for both parameters, within the following bounds: 0<fc_init_tot<2; 0<fc_processing<15. (B,D,F,H) Model fits using the mean values of the posterior sample distributions for the two parameters.

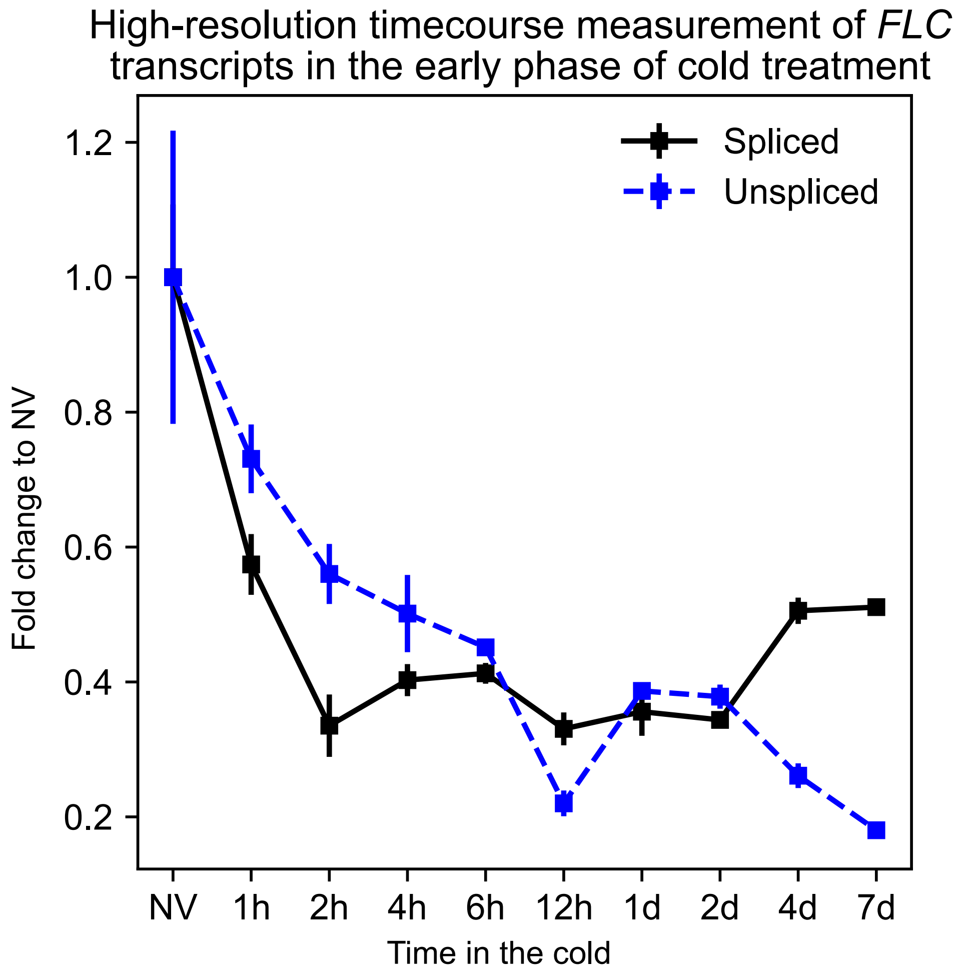


**Figure S12. Fold changes in cold induced reduction of spliced and unspliced *FLC* transcripts show opposite trends over short and long timescales**. Relative expression of spliced and unspliced *FLC* (as defined in the Results section – also see list of primers) over a vernalization time course (constant 5°C) in Col*FRI*. Values are normalized to the housekeeping gene UBC and to the NV levels. Error bars represent SEM (n=4 for NV timepoint, n=3 for all other timepoints). Over the first 6h of cold treatment, spliced *FLC* level reduces faster than unspliced *FLC.* This trend is reversed over the 2 to 7 day phase, where unspliced *FLC* level reduces and spliced *FLC* level shows no further reduction. This is consistent with the long term trend observed in Fig. 1(A), where the relative reduction in unspliced *FLC* is significantly greater than for spliced *FLC* over 2, 4, 6 and 8 week timepoints.

### Table S1

#### Parameter values for *FLC* intron 1 processing model

Timepoints

| | Transcription initiation rate | Pol II speed | Intron processing rate | 5'-3' degradation rate | $f_{lariat}$ |
|---|---|---|---|---|---|
| **NV** | - | 0.04kb/s | 0.002/s | 0.003kb/s | Fitted |
| **2WT0** | Fold change inferred | Fold change 0.8 | Fold change inferred | 0.003kb/s | Fitted |
| **4WT0** | Fold change inferred | Fold change 0.65 | Fold change inferred | 0.003kb/s | 0.17 |
| **6WT0** | Fold change inferred | Fold change 0.6 | Fold change inferred | 0.003kb/s | 0.17 |
| **8WT0** | Fold change inferred | Fold change 0.6 | Fold change inferred | 0.003kb/s | 0.17 |
| **2WT7** | Fold change inferred | Fold change 0.6 | Fold change inferred | 0.003kb/s | 0.17 |
| **4WT7** | Fold change inferred | Fold change 0.5 | Fold change inferred | 0.003kb/s | 0.17 |
| **6WT7** | Fold change inferred | Fold change 0.3 | Fold change inferred | 0.003kb/s | 0.17 |
| **8WT7** | Fold change inferred | Fold change 0.3 | Fold change inferred | 0.003kb/s | 0.17 |

**Table S1. Parameter table for the *FLC* intron 1 processing model, showing values used for fixed parameters and indicating which parameters were inferred at each timepoint.**
